# Bridging the gaps between field-based ecology and remote sensing to estimate plant functional diversity: a systematic review

**DOI:** 10.64898/2026.02.19.706857

**Authors:** José M. Cerda-Paredes, Javier Pacheco-Labrador, Dylan Craven, Javier Lopatin

**Affiliations:** Data Observatory, Santiago, Chile; Facultad de Ingeniería y Ciencias, Universidad Adolfo Ibáñez, Santiago, Chile; Environmental Remote Sensing and Spectroscopy Laboratory (SpecLab), Spanish National Research Council, Madrid, Spain; GEMA Center for Genomics, Ecology & Environment, Universidad Mayor, Santiago, Chile; Center for Climate Resilience Research (CR)2, University of Chile, Santiago, Chile

**Author notes:** Joint senior authors (Dylan Craven, Javier Lopatin).

## Abstract

Understanding plant functional diversity across scales requires integrating field-based ecology and remote sensing, yet these disciplines differ in how traits are studied. We evaluated the conceptual and methodological convergence between these disciplines. Our results reveal that field-based ecology has undergone longer conceptual development and covers a broader range of traits, while remote sensing has experienced rapid growth driven by technological advances. Both disciplines are increasingly converging on similar concepts. However, major gaps in empirical coverage persist across biomes in both disciplines. Although plant-dominated ecosystems have been extensively studied, extreme ecosystems remain undersampled. While there is considerable diversity in the definition “functional traits”, both disciplines converge on using a similar set of traits, reflecting their central role in plant strategies and spectral detectability. Our synthesis underscores the potential for methodological synergy. Harmonizing trait definitions, scaling assumptions, and computational steps involved in estimating plant functional diversity are crucial for building a unified, multiscale framework for biodiversity monitoring in ecosystems undergoing biodiversity loss and climate change.

**Teaser:** A synthesis of how field ecology and remote sensing can be aligned to monitor plant functional diversity across scales.

## Introduction

The current biodiversity crisis remains inadequately addressed due to significant gaps and biases (e.g., taxonomic and spatial) in biodiversity data (*1, 2*). Despite high biodiversity loss rates, we still lack a clear understanding of the when, where, how, and why of biodiversity change, and hence its implications for vital ecosystem services (*3*). A key step towards closing these knowledge gaps is the establishment of a global monitoring system that systematically monitors multiple facets of biodiversity (i.e., taxonomic, phylogenetic, and functional diversity) and ecosystem services from local to global scales (*4*). One of the ways this need has been addressed is through the Essential Biodiversity Variables (EBVs) framework (*5*), which is designed to standardize global biodiversity monitoring across ecological and spatial scales. Several EBVs – such as species traits and community composition– explicitly include components of functional diversity that remote sensing is expected to quantify (*6*). In remote sensing, this contribution usually works by estimating individual plant traits, from which functional diversity can be calculated although alternative approaches may bypass explicit trait estimation. Among biodiversity facets, functional diversity has become essential for monitoring plant responses to climate and land-use change (*7*) and identifying regime shifts in ecosystems (*8*). However, cross-scale monitoring of functional diversity relies upon approaches –principally field-based ecology and remote sensing– that seek to measure the same entities despite employing what at first glance appear to be contrasting concepts and methods.

Traditionally, functional diversity describes the distribution of functional traits (i.e., any trait that influences the fitness of an individual organism) in an assemblage (*9–11*). However, substantial debate remains regarding which traits should be considered functional, how to weigh them, and which metrics best capture different aspects of functional diversity (*10*, *12*, *13*). In practice, functional diversity is commonly interpreted more broadly as the variation or distribution of traits within communities, including those related to ecological strategies or ecosystem functions, regardless of whether they directly impact fitness or are measured at the individual scale. Functional diversity can be decomposed into functional richness, evenness and divergence. These components of functional diversity identify the range of ecological strategies present (i.e., richness), how uniformly traits are distributed (i.e., evenness), and how much co-occurring species differ functionally (i.e., divergence) (*13–15*). In addition, other multidimensional indices such as functional dispersion (*14*) or Rao’s quadratic entropy (*Q*) (*16*) are widely used to quantify trait dissimilarity within communities and are often interpreted as capturing functional divergence. Furthermore, integrative frameworks have been proposed to relate richness, evenness, and divergence within a unified diversity metric [e.g., (*17*, *18*)]. Additionally, community-weighted trait means (CWM) express the trait values of dominant species in an assemblage (i.e., functional composition), and are therefore expected to have a direct effect on ecosystem functioning (“mass ratio hypothesis”) (*19*), as are other statistical moments of individual trait distributions [e.g., variance, skewness, and kurtosis (*20*)]. The variety of components and metrics used to quantify functional diversity allows for considerable flexibility in cross-scale biodiversity monitoring. However, it also influences how different disciplines implement these approaches. Not all functional diversity metrics are equally transferable or comparable between field-based and remotely sensed data (*21–23*).

The different methods and technologies used by field-based ecology (i.e., direct measurement and destructive or non-destructive sampling of plant tissues) and remote sensing (i.e., spectral imagery, imaging spectroscopy, multispectral reflectance, LiDAR and radar) translate into differences in the spatial, temporal, and ecological scales of plant functional diversity. In this context, we use *field-based ecology* to refer to studies that use empirical trait data collected in natural, semi-natural and experimental ecosystems. In terms of spatial scale, remote sensing approaches estimate functional diversity at the spatial resolution of the sensor [i.e., pixel size; (*24*)]. On the other hand, field ecology approaches measure traits at the individual, species, or community levels (*25*), and is often done so at resolutions smaller than pixels. A second key difference between approaches are the sampled individuals selected for estimation functional diversity. Field ecology computes diversity within comparable levels of organization (e.g., traits of individuals belonging to the same community). Remote sensing typically estimates functional diversity by comparing spectral across spatial elements (i.e., among neighboring pixels or within ecologically meaningful units) (*25–28*). In practice, these comparisons are commonly implemented using spatial aggregation schemes, such as moving-window approaches or predefined vegetation units.

Beyond spatial scale, remote sensing and field-based ecology also differ in the type of variables they emphasize, as remote sensing studies often estimated functional diversity directly from spectral properties or plant biophysical properties based on spectral properties (*22*, *29*), some of which can be considered plant functional. Meanwhile, field-based ecology typically quantifies functional diversity from morphological, physiological, or biochemical traits measured *in situ* or with trait values from local, regional, or global databases. Remote sensing can also use field data for validation of plant trait estimates [e.g., (*21*)], but these do not necessarily match those sampled in field ecology since they prioritize those that exhibit strong correlations with spectral signals. Consequently, the traits or trait proxies representing plant function may reveal different facets of plant form and function, and therefore are not always comparable between the two approaches (*12*, *22*). In terms of temporal scales, functional diversity can be estimated frequently and continuously with remote sensing products, because satellite sensors revisit locations at regular intervals, thereby providing opportunities to capture seasonal or interannual dynamics (*30–32*), whereas field-based estimates of functional diversity are usually limited to discrete sampling campaigns, providing high-resolution but temporally localized snapshots of trait distributions and diversity (*10*, *33*).

Remote sensing-based estimates of functional diversity generally represent the community level, although this is not always stated explicitly [but see (*23*)]. Field-based approaches, in turn, estimate functional diversity across a range of ecological levels, from individuals to communities, by scaling up trait measurements using species abundances or biomass (*19*, *34–41*). Given the logistical constraints associated with trait measurements (*10*), field-based trait measurements rarely cover all individuals in a community or population. Despite their differences, both approaches employ similar analytical tools. For example, both often reduce the dimensionality of trait or spectral data using principal components analysis (PCA) or principal coordinates analysis (PCoA), and estimate some or all components of functional diversity (*13*). Taken together, these methodological differences reflect scale-dependent trade-offs rather than contradictions, and highlight how each discipline captures complementary dimensions of functional diversity (*22*).

Despite their shared goal of quantifying functional diversity, the methodological differences between field-based and remote sensing approaches have not been systematically evaluated, including how plant functional diversity is conceptualized, measured, and applied across these two disciplines, leaving key challenges and possible solutions for integration largely unexplored. Reviews of field-based functional diversity have primarily focused on how plant traits contribute to resource use and ecological performance, effects of spatial and temporal scales, and the quantification of trait variability (*15*, *33*, *42*). Meanwhile, remote sensing reviews have identified the methodological challenges of mapping plant traits (but not diversity), principally data gaps due to taxonomic, trait, and spatial biases or multi-sensor data complementarity (*43*, *44*), and only more recently, methodological developments to infer plant diversity (*28*, *45*). Linking plant traits measured *in situ* with remote sensing data continues to be a common challenge. While some reviews have proposed strategies for upscaling field-based trait data with remote sensing (*46*, *47*), they also note that environmental conditions (e.g., illumination, canopy structure, or soil background), as well as changes in these conditions induced by land-use or vegetation disturbances can alter canopy reflectance and thereby influence the accuracy and transferability of remotely-sensed estimates of plant functional diversity. This underscores the importance of combining *in situ* trait data with remote sensing to improve the robustness and ecological interpretability of functional diversity estimates across data sources, ecosystems, and spatial resolutions (*12*).

Here, we aim to identify the methodological and conceptual challenges of developing a cross-scale approach to quantifying and monitoring plant functional diversity. To do so, we evaluate how key concepts and methods are used in field-based ecology and remote sensing approaches to quantifying plant functional diversity by combining a bibliometric analysis with a systematic review (“research weaving”) (*48*). First, we analyze the development of plant functional diversity research in field-based ecology –studies measuring in natural, semi-natural, and experimental ecosystems– and remote sensing. We then systematically review the conceptual and methodological similarities and differences in spatial scales (i.e., resolution and extent), ecological characteristics and scales (i.e., traits, biomes, ecological scale), and functional diversity indices used in both approaches.

## Results & Discussion

Our bibliometric analysis reveals differences in the historical trajectories and geographical patterns of research about plant functional diversity in field-based ecology and remote sensing. Field-based ecology has a longer publication history and greater conceptual maturity in plant functional diversity (*K* < 0.75; Fig. 1A and Table S2). Consequently, field-based ecology has developed more consolidated theoretical frameworks and well-established methodologies (*10*, *11*, *13*, *15*, *33*, *39*). In contrast, remote sensing is undergoing rapid growth and remains in the *revolution stage* (*K* > 0.75; Fig. 1A and Table S2), as the process of adapting and validating conceptual frameworks for functional diversity is ongoing and fragmented (*22*, *26*, *43*, *44*, *49–52*). *K* also changes more slowly for remote sensing (slope = −0.011) than for field-based ecology (slope = −0.013) over time, which could be due to the fact the discipline is still constrained by current technological limitations, the capacity to measure certain plant traits (e.g., root traits), and challenges integrating ecological theory and *in situ* data (*53*). Across both disciplines, these historical differences are accompanied by pronounced geographical biases in research effort and data availability. Global plant trait databases remain strongly skewed towards the Global North, with major gaps across tropical, arid, and high-mountain ecosystems (*54*) Consistent with this pattern, our results show that the growing body of research on plant functional diversity exhibits marked differences in scientific productivity (i.e., number of articles published) between the Global North and South. Although both disciplines exhibit a pronounced bias towards the Global North, this pattern is more pronounced in field-based ecology, where research activity is concentrated in fewer countries than in remote sensing (Fig. S3).

**Fig. 1.**
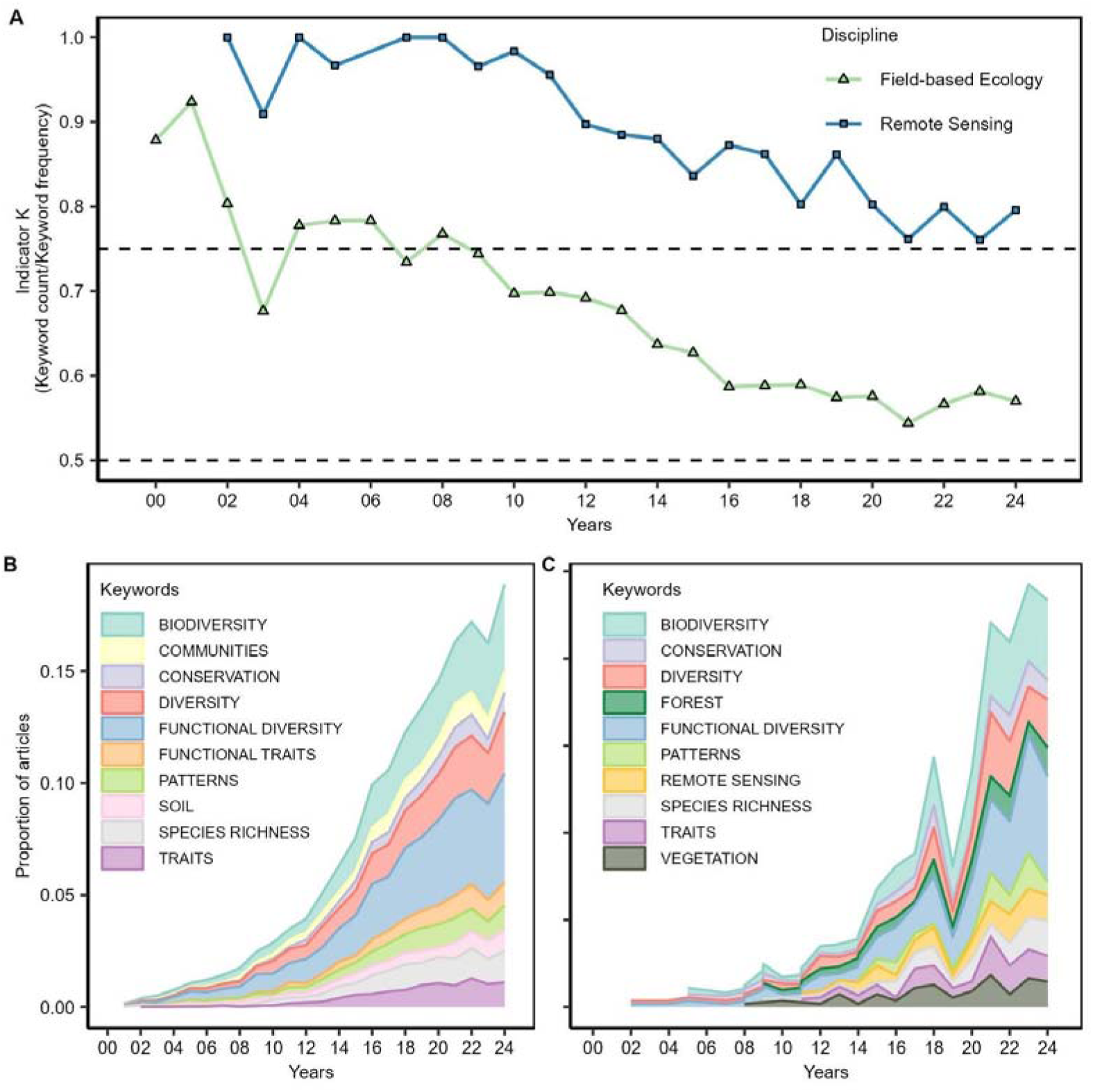
Temporal evolution of plant functional diversity research. (**A**) Temporal evolution of indicator K for scientific literature about field-based ecology and remote sensing approaches to quantifying functional diversity. The indicator K quantifies the maturity of a scientific discipline, ranging from a pre-revolution or revolution stage (1 - 0.75) to a forming or pre-normal science stage (0.75 - 0.5) (*144*). (**B)** Temporal evolution of top 10 most frequently used keywords in field-based ecology literature. (**C**) Temporal evolution of top 10 most frequently used keywords in remote sensing literature. Circles indicate keywords used in both disciplines, squares indicate keywords exclusive to remote sensing, and triangles indicate keywords exclusive to field-based ecology.

Patterns in keyword frequency point to an increasing overlap in thematic emphasis between field-based and remote sensing studies. We thus observed growing conceptual convergence between the two disciplines (Fig. 1B and C; Fig. S4 and S5). For example, we found that “functional diversity” and “biodiversity” were among the most frequently used keywords in both disciplines (Fig. 1B and C; Fig. S4 and S5), reflecting shared interests even if the terms may not be used in conceptually equivalent ways. This could indicate that field ecologists are increasingly adopting remote sensing tools at the same as remote sensing researchers are incorporating conceptual frameworks focused on functional diversity into their research. This integration is consistent with the growing emphasis on interoperability across scales and disciplines within global biodiversity monitoring initiatives (e.g., GEO BON, Biodiversity+ Precursors and EO4Diversity). In this context, EBVs such as species traits and community composition explicitly include traits, which remote sensing is expected to be able to estimate (*5*, *6*, *55*).

Our structural topic model identified seven key topics in the literature on plant functional diversity and their temporal evolution over the past three decades (Fig. 2). The most dominant topic in this body of literature is the identification of the environmental drivers of trait diversity (Topic 2; Fig. 2A). Studies that address the effects of environmental gradients (e.g., temperature, precipitation, elevation, and land-use change) tend to be more generalizable and comparable across regions, ecosystems and scales (*56*). We found that there was general agreement on most of the main topics between disciplines, as the probability of most topics did not vary significantly (P > 0.05; e.g., Topic 1, 2, 4 and 6; Fig. 2B). However, we observed a pronounced lag in the evolution of key topics between disciplines: the proportions of topics stabilized in field-based ecology around 2004, but not until 2015 in remote sensing (Fig. 2C and D). Beyond temporal dynamics, topics encompassing belowground ecology, such as root traits or microbiological communities are mostly addressed by field ecologists (P < 0.05; Topic 3 and 7; Fig. 2B). Topics related to belowground ecology are particularly challenging to quantify with remote sensing, which has limited their use in global or comparative studies (*57*). Although a few attempts have been made to estimate belowground traits using remote sensing (e.g., SAR- or model-based approaches), these remain rare and not widely applicable across ecosystems (*58*, *59*). Remote sensing literature is more strongly associated with Topic 5, which represents general aspects of research practice, than field-based ecology (P < 0.05; Topic 5; Fig. 2B) possibly because it is still in the revolutionary stage as a discipline, and this topic can be effectively addressed using information derived from remotely sensed plant data. This suggests that remote sensing still requires further development of methodological frameworks to improve the ecological validity and cross-scale comparability of remotely-sensed functional diversity estimates. Additionally, there needs to be further integration of conceptual frameworks from field-based ecology, which has already led to conceptual convergence and methodological divergence between the two disciplines (Fig. 1A). Also, as remote sensing is inherently limited by its detection capability, topics associated with species-level patterns are significantly associated with field-based ecology (P < 0.05; Topics 3 and 7; Fig. 2B).

**Fig. 2.**
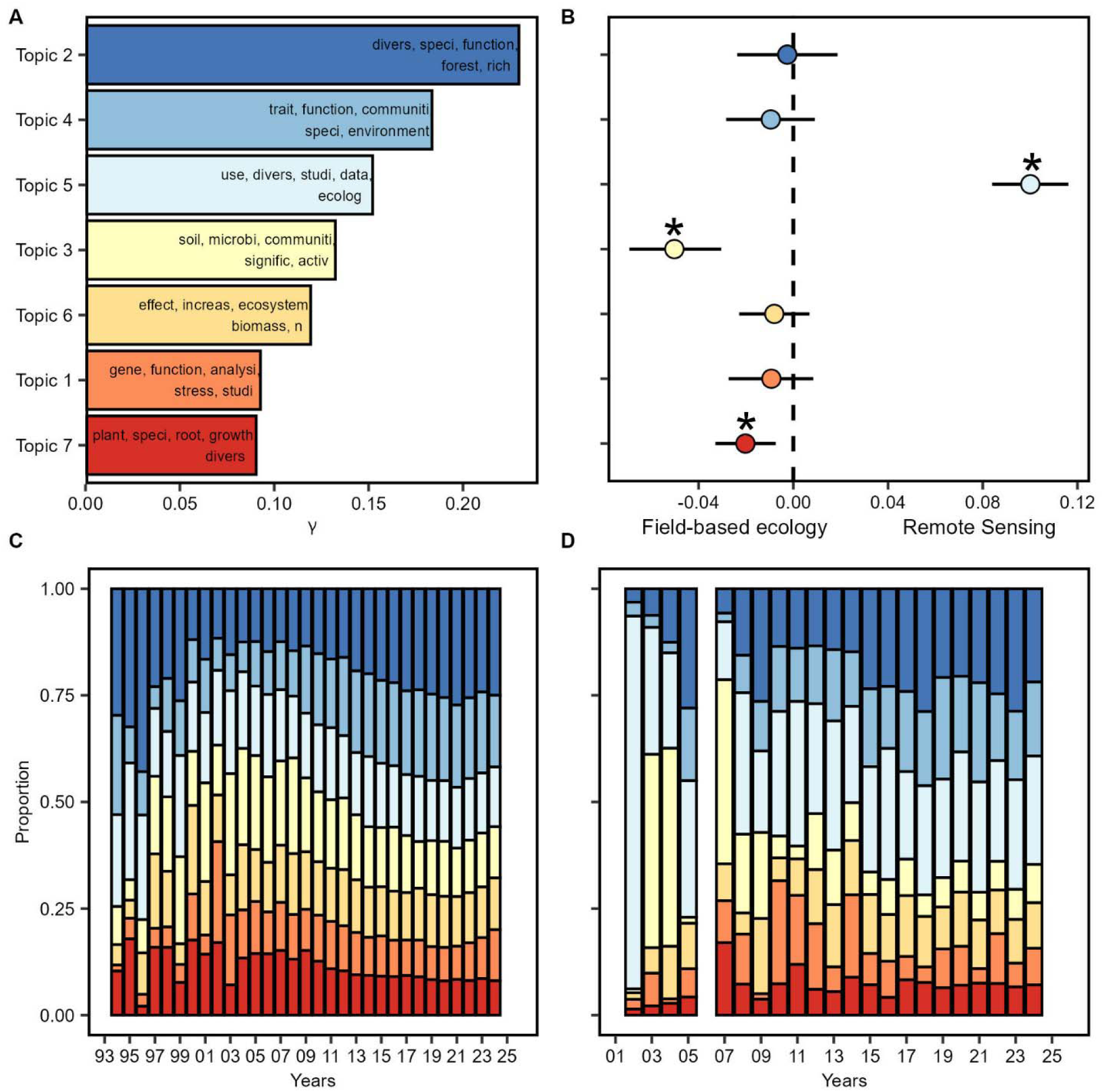
Temporal evolution of key topics in scientific literature to quantifying plant functional diversity. (**A**) Seven topics identified by a structural topic model (STM), ranked by prevalence and the top 5 terms of each topic. (**B**) Effect of discipline on topic prevalence. (**C**) Estimated proportion of each topic annually in field-based ecology. (**D**) Proportion of each topic annually in remote sensing. γ: mean probability of each topic that an article belongs to. Significant differences in covariance (P < 0.05) are denoted by asterisks.

### Ecological dimensions of plant functional diversity research

Our results reveal an asymmetric distribution of research on plant functional diversity across biomes. We identified plant functional diversity studies conducted across 13 biomes, but found that remote sensing has assessed plant functional diversity patterns in only seven of them, mainly those dominated by forests, savannas, and grasslands (Fig. 3). The limited research on the land-water interface (e.g., palustrine wetlands, shorelines, lakes) and extreme biomes (e.g., deserts and semi-deserts) highlights a major gap in our understanding of plant functional diversity. These biomes have been studied mainly using fieldwork-based approaches, despite logistical limitations such as accessibility and high fieldwork costs. In land-water interface ecosystems, remotely sensed detection of plant functional traits can be challenging because vegetation often occurs in small patches, and the presence of water can hinder the interpretation of spectral signals (*60*, *61*).

**Fig. 3.**
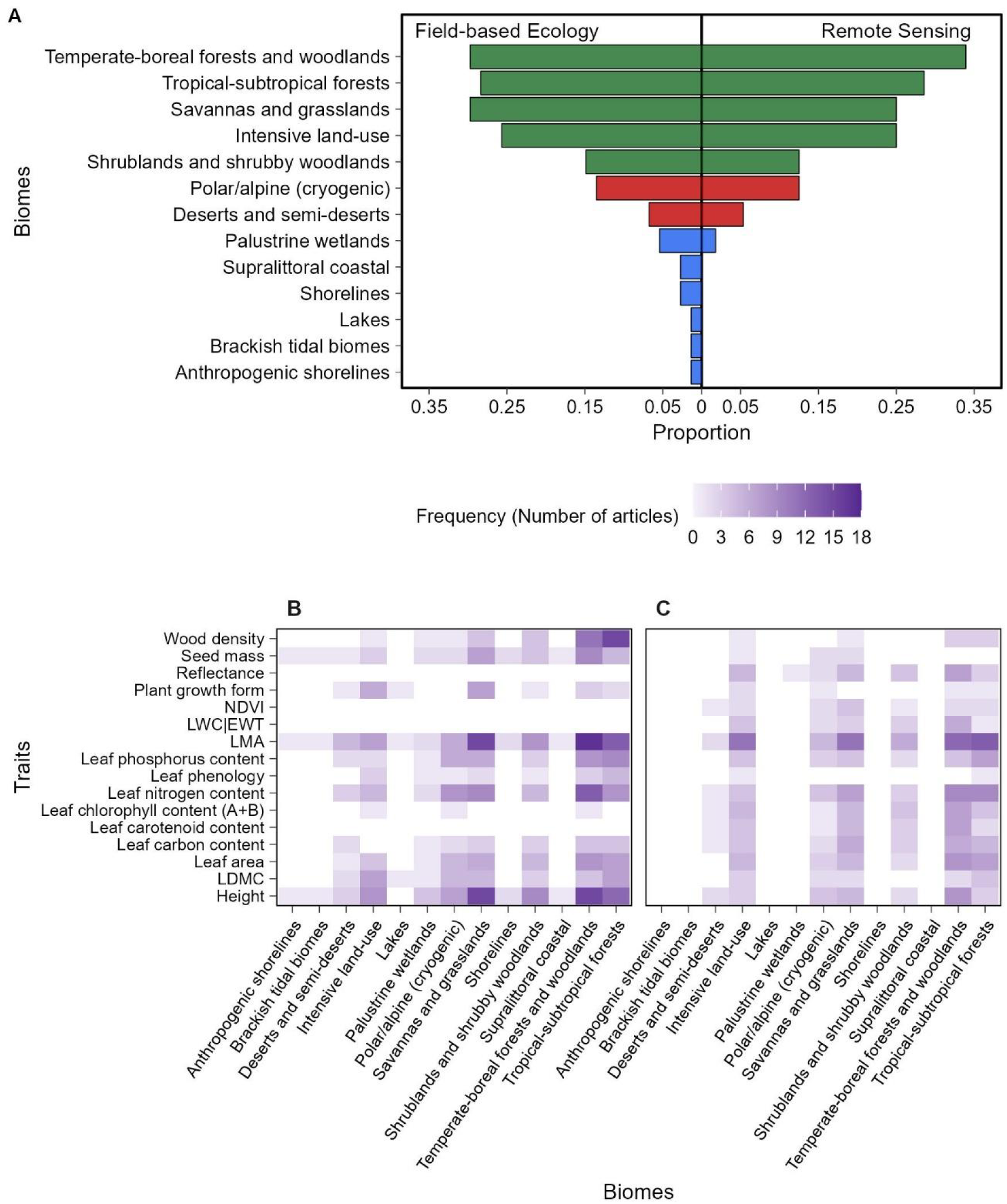
Biome and trait coverage in functional diversity research. (**A**) Proportion of reviewed articles in each biome, relative to the total number of articles analyzed for each discipline. (**B**) Functional traits assessed in each biome in field-based ecology. (**C**) Functional traits evaluated in each biome in remote sensing. In (**A**) bar colors represent categories of biomes, blue are land-water interface biomes, red are extreme biomes, gray are intensive land-use biomes and green are highly dominated by vegetation biomes. We included functional traits with an overall frequency ≥ 10. NDVI: Normalized difference vegetation index (*150*), LWC: Leaf water content, EWT: Equivalent water thickness, LMA: Leaf matter per area, LDMC: Leaf dry matter content.

In desert ecosystems, the low vegetation cover and the high albedo of soils and sand obscures the signal of vegetation, often saturating optical bands (*62*). Similar to land-water interface ecosystems, scarce but clumped vegetation in deserts reduces the reliability of medium and low resolution remote sensing outputs and complicates trait-based analyses (*63*, *64*). Nevertheless, some functional traits can still be obtained with remote-sensing products in these ecosystems, such as pigment content (e.g., chlorophyll) or leaf water content in land-water interface ecosystems [e.g., (*65*, *66*)], and indicators of physiological stress in deserts, such as pigment ratios [e.g., (*67*)]. Given the increasing ecological importance of these underrepresented ecosystems under climate change scenarios, expanding functional diversity research into less studied biomes is essential for a comprehensive and balanced understanding of biodiversity (*68*, *69*). However, the current limitations in spatial resolution frequently restrict functional diversity studies to ecosystems dominated by large plants (e.g., forests) or to homogeneous low-height ecosystems (e.g., grasslands). Although functional diversity is widely studied in both biomes using remote sensing, it is not always successful (*70*).

Beyond geographical biases in plant functional diversity, analysis reveals notable differences in how each discipline selects, defines, and uses plant traits. We identified an extensive list of traits used for estimating plant functional diversity (140 plant traits; Fig. S6). Rather than reflecting a disagreement between disciplines, this diversity primarily arises from the broad range of trait types commonly employed in functional ecology. These trait types span fitness-related traits, response and effect traits, and easily measured morphological or physiological traits (*9*, *11*). Since the term “functional trait” is interpreted flexibly in ecological studies, researchers often select traits based on their goals, ecological context, or data availability. Similarly, remote sensing studies distinguish between spectral, structural, and physiological proxies and biophysical traits (*12*, *53*), while also inheriting part of the conceptual breadth present in trait-based ecology (*51*). Despite this diversity of approaches, both disciplines converge on a shared subset of commonly used traits (Figs. 3 and 4).

**Fig. 4.**
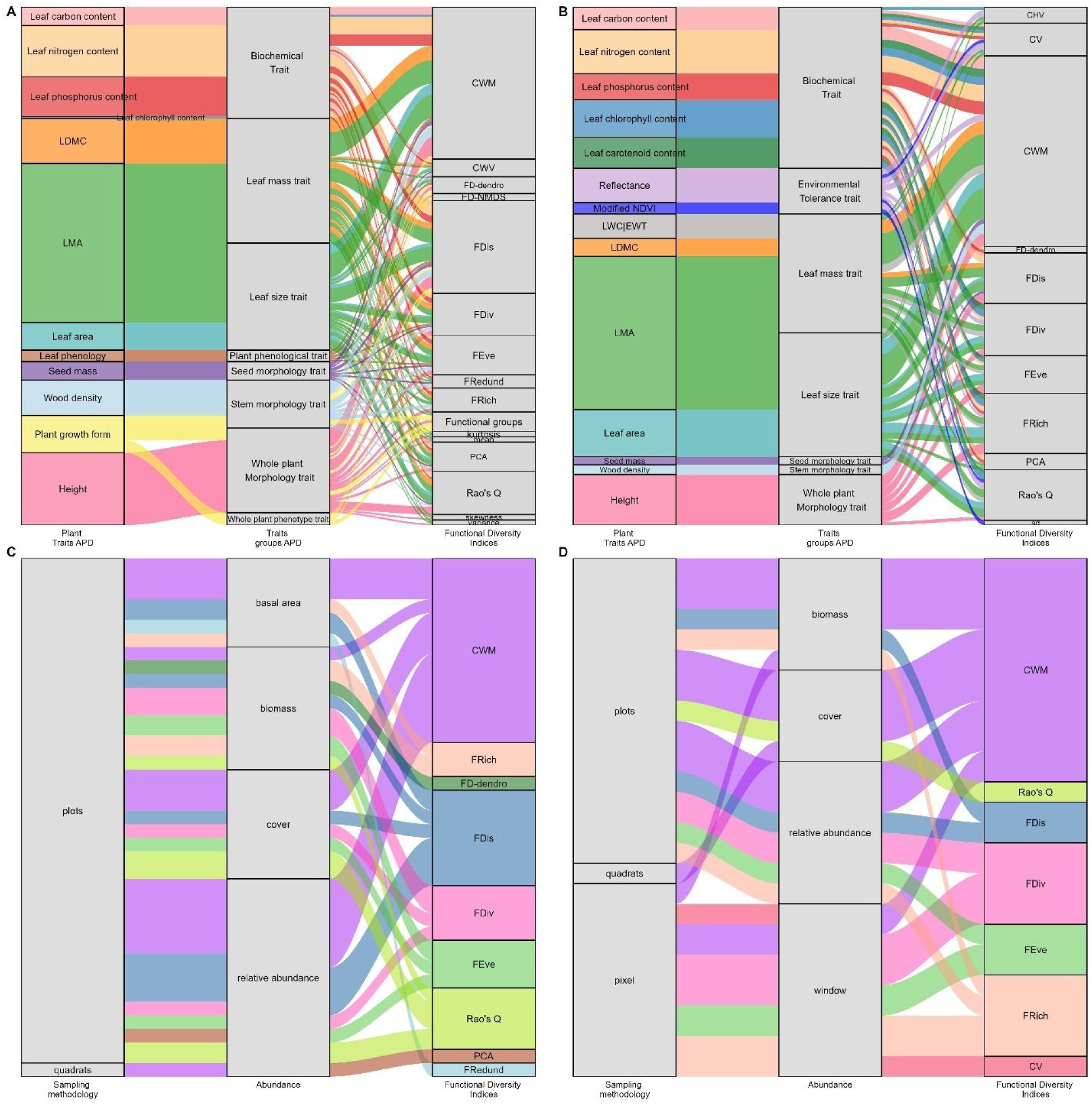
Linking traits, abundance, and functional diversity across disciplines. **(A)** and **(B)** connect plant traits, trait groups, and functional diversity indices for (**A**) field-based ecology and (**B**) remote sensing. (**C**) and (**D**) connect sampling methodology, abundance variable and functional diversity indices for (**C**) field-based ecology, and (**D**) remote sensing. In (**A**) and (**B**) we included functional traits with an overall frequency ≥ 10 and functional diversity indices with at least two records per trait. In (**C**) and (**D**) we included functional diversity indices with at least two records per abundance variable. APD: AusTraits Plant Dictionary (*147*), LDMC: Leaf dry matter content, LMA: Leaf matter per area, NDVI: Normalized difference vegetation index (*150*), CWM: Community weighted mean, CWV: Community weighted variance, FD-dendro: Functional diversity as sum of branch length of classification, FD-NMDS: Functional diversity from non-metric multidimensional scaling, FDis: Functional dispersion, FDiv: Functional divergence, FEve: Functional evenness, FRedund: Functional redundancy, FRich: Functional richness, PCA: Principal component analysis, LWC: Leaf water content, EWT: Equivalent water thickness, CHV: Convex hull volume, CV: Coefficient of variation, sd: Standard deviation.

Both disciplines also diverge substantially in how trait data are collected and upscaled. Field-based ecology typically relies on direct trait measurements from individuals or populations, providing high ecological resolution, but limited resolution and temporal coverage (*10*). In contrast, remote sensing often uses spectral proxies instead of direct measurements [e.g., (*71*, *72*)], enabling broader spatial coverage but introducing uncertainty because spectral signals often integrate information from multiple individuals or species within a pixel. These differences in trait acquisition and scaling influence how traits are represented in functional diversity indices and ultimately affect the comparability of estimates between disciplines.

Trait frequency trends indicate that only a few traits are consistently represented in the literature of both disciplines (Fig. 3B and C). Among the traits most frequently used in both field-based and remote sensing studies are plant height (or canopy height), leaf mass per area (LMA), and leaf nitrogen content (N; Fig. 3 and 4). Leaf nitrogen is reported in the literature as either mass-based concentration or area-based content. However, from a radiative transfer perspective, area-based expressions are more directly linked to canopy reflectance. Concentration-based estimates, on the other hand, may largely reflect variation in leaf mass per area (*73*). This can limit their interpretability when derived from remote sensing (*74*). These traits are commonly used to upscale from the individual to the canopy levels (*75*, *76*). Although categorized as soft traits, they can also be considered context dependent, as their ecological relevance and predictive capacity vary across environmental contexts and/or biological interactions (*77–80*). For instance, plant height is central to vegetation structure and competitive dynamics for light, operating through frequency-dependent interactions (*81*, *82*). LMA (or its inverse, SLA) and leaf nitrogen content are integral components of the leaf economics spectrum rather than physiological traits in the strict sense, representing fundamental trade-offs between resource acquisition and conservation (*77*, *83*). Studies have shown that these leaf economic strategies can also be detected through spectral information (*74*, *84*). This strong link between leaf economic traits and whole-plant strategies explains why these traits are a primary focus in field-based studies.

LMA and plant height are relevant in remote sensing as they directly influence vegetation spectral properties and vertical structure, both of which can be remotely observed. For example, canopy height can be directly observed with LiDAR data (e.g., GEDI) or estimated via photogrammetry and interferometry processes (e.g., drone or Sentinel-1), providing high-resolution vertical structural data. Meanwhile, LMA directly influences leaf optical properties, mostly in the short-wave infrared region (*85*), and can be estimated via empirical or radiative transfer models (RTM) [e.g., (*84*)]. The frequent use of LMA and plant height in both disciplines aligns with the foundational L-H-S strategy (*86*), which identifies these as core traits that capture key ecological trade-offs, such as resource use (i.e., LMA) and competition for light (i.e., height). This convergence suggests that despite methodological differences, both disciplines prioritize traits central to global plant strategy spectra (*7*, *81*, *87*). On the other hand, seed and root traits are underrepresented in remote sensing literature (Fig. 3 and 4), highlighting a significant technological limitation in capturing reproductive and belowground strategies (*88*), which are often harder to observe remotely (*44*) despite their central role as response and effect traits that mediate ecosystem sensitivity to environmental change (*40*, *89*).

The number and types of traits measured by each discipline show considerable differences. Field-based ecology encompasses a broader array of plant traits than remote sensing (n = 122 and n = 84, respectively; Fig. S7). Traits such as wood density (dry matter per volume) and leaf dry matter content (LDMC; dry mass per fresh mass), frequently used in field-based studies, are also context dependent and relate to plant investment in structural and physiological strategies (*90–92*). For example, LDMC provides insight into plant strategies related to resource conservation, structural investment, and adaptation to environmental stress (*80*). Interestingly, remote sensing studies often use leaf water content (LWC; mass water per area; Fig. 3 and 4), to assess similar physiological properties as LDMC, such as water retention or tissue investment. Remote sensing studies often use area-related traits (contents; mass per area), which interact with light during the radiative transfer process, whereas field-based ecology commonly expresses these same variables as concentrations (mass per mass), reflecting compositional rather than radiative properties. Remote sensing studies also sometimes use concentration for depicting pigments in the reflectance signal, but this has shown to be heavily related more to leaf mass per area than to pigments itselves (*74*). However, field-based ecology often uses mass-based trait values; mass-and area-based trait values capture different components of leaf structure and physiology that can influence ecological interpretation (*93*). These estimations do not require optical modeling or spectral calibration. This distinction does not imply a disadvantage for either approach, but rather reflects the fact that radiative transfer models strictly represent leaf properties as contents. This means that variables such as pigments, water, and dry matter are inherently expressed in these units (*94–96*). These different measurement conventions stem from the methodological principles of each discipline. While differing in methods and scales, both disciplines aim to capture mechanistic aspects of plant performance, using distinct –but possibly complementary– traits. The empirical and biophysical links between plant traits and spectral reflectance offer significant potential to remotely infer plant function, enabling large-scale assessments of plant strategies and ecosystem functioning (*84*, *97*).

Patterns in trait detectability reveal that remote sensing studies tend to focus on a narrower subset of traits shaped by optical and biophysical constraints. Remote sensing emphasizes biochemical traits (e.g., leaf carbon, phosphorus, and chlorophyll content) and spectral traits (e.g., reflectance factors and spectral indices) linked to the interaction of light and matter [see (*98–100*)], and encompass the variability of multiple traits [or properties that might not be considered traits *sensu stricto* (*11*)], but also confounding factors, such as bare soil or shadows (Fig. 3B and C). Pigments, water and dry matter traits are often included as parameters of the most used radiative transfer models such as PROSPECT (*94*) or FLUSPECT (*95*). The selection of traits in remote sensing studies is strongly influenced by their detectability at the canopy scale, limiting the inclusion of traits related to reproduction, growth form, belowground strategies, phenology, or biotic interactions [e.g., seed mass, flower height and diameter, root traits, flowering time, nitrogen fixation, or mycorrhizal associations; (*101–103*)]. These traits are still predominantly measured using field-based approaches, although remote sensing continues to expand the range of traits for which meaningful proxies can be derived. While efforts to estimate some of these traits using remote sensing are ongoing [e.g., (*104*)], their limited abundance and representativeness in canopy reflectance, or their temporal variability, could present technical challenges. The spatial heterogeneity introduced by human-modified landscapes exacerbates trait heterogeneity, where regional context (i.e., species introductions, land-use history, and novel communities) can alter or weaken trait-environment relationships (*47*, *105*). This added variability increases uncertainty in the estimation of trait-related spectral signals [e.g., (*106*)], which in turn affects the robustness of remote sensing upscaling.

Our analysis also reveals that remote sensing uses response traits to a greater extent than field-based ecology (*40*, *89*, *107*). Whereas remote sensing often captures response traits linked to environmental gradients (*105*), it also commonly retrieves canopy-level properties associated with ecosystem functioning (e.g., productivity proxies such as NDVI). In contrast, field-based ecology more commonly includes structural and reproductive traits linked to environmental responses and ecosystem effects (*108*). However, these tendencies are not absolute, and both disciplines are increasingly expanding the range of traits and processes they can address. Together, these disciplines provide complementary insights into how plants respond to and influence their environments. At the same time, neither discipline alone captures the full range of organismal strategies and ecosystem functioning. Bridging both disciplines will therefore be essential for monitoring biodiversity responses to global change and how these responses may affect ecosystem functioning.

### Functional diversity indices

Despite differences in trait selection, functional diversity indices showed clear convergence across disciplines (Fig. 4). Nearly half of field-based studies (47%) and more than one-third of remote sensing studies (37%) used CWMs (Fig. 4), and functional dispersion was also widely applied in both disciplines (Fig. 4A and B), highlighting a shared conceptual foundation in quantifying functional diversity. In addition, field-based studies tend to use indices that capture multiple components of community structure (i.e., functional dispersion, evenness, and divergence; Fig. 4A and C), while remote sensing studies emphasize particular aspects of community structure (i.e., CWM and functional richness; Fig. 4B and D). This divergence largely reflects not only methodological constraints and data properties (e.g., differential sensitivity to data dimensionality, trait covariance structure, and spectral aggregation), but also conceptual gaps in remote sensing. The ecological interpretation of trait distribution remains underdeveloped. For example, some indices, such as functional divergence and Rao’s Q may retain a stronger ecological signal of functional differentiation, whereas evenness-based metrics can be more sensitive to methodological choices and aggregation effects (*22*). These differences affect how each discipline evaluates patterns and drivers of functional diversity. Remote sensing often highlights shifts in dominant ecological strategies with CWM-based metrics, whereas field-based studies emphasize trait variation. However, in remote sensing, CWM-based metrics usually reflect mean trait values derived from pixels or aggregated canopy signals, not abundance-weighted species means. This distinction reflects the difference between biological and radiative representations of community composition. Together, these perspectives offer complementary insights into how communities respond to environmental gradients. Yet, interpreting functional diversity is inherently scale dependent in both disciplines (*109*). In remote sensing, this context dependency is further influenced by pixel-level variability, where spectral signals often integrate information from mixed plant communities. Consequently, the ecological meaning of spectral or trait-based diversity may differ from species- or community-level definitions (*110*).

Our results show that abundance is incorporated through different methodological strategies, each contributing complementary insights into community-level functional diversity (Fig. 4C and D). Both disciplines use abundance-weighted metrics to upscale trait diversity to the community level, giving more importance to species with higher dominance. However, both disciplines often use different methodological approaches (Fig. 4C and D). Field-based studies derive these weights directly from *in situ* measurements of biomass, cover, and relative abundance.

Whereas methodological uncertainties affect both disciplines, differences in how spatial scale is inherently represented in field-based and remote sensing functional diversity indices introduce specific challenges for cross-disciplinary applications. Field-based studies most commonly quantify α-functional diversity at the plot level, but can also be extended to assess β- and γ-functional diversity across multiple plots (or sites). These field-based studies often rely on spatially discrete sampling designs across discontinuous spatial extents (*21*, *111*). In contrast, remote sensing can capture broader-scale patterns (β and γ) by quantifying α-functional diversity across continuous spatial extents, or by deriving γ from additive (α + β) or multiplicative (α × β) partitioning frameworks (*50*). In remote sensing, these frameworks can be extended through Rao’s Q-based measures of spatial and temporal diversity (*25*), further demonstrating how can shape the comparability between disciplines. This distinction is essential because comparing field-based and remotely-sensing functional diversity requires aligning trait definitions, metrics, and the scale. Notably, spatial extent is not explicitly reported in 81% of the field-based and 55% of remote studies we reviewed, further complicating cross-study comparisons. Without explicitly aligning the spatial scales of assessment, discrepancies in functional diversity values may arise from scale mismatches rather than intrinsic methodological performance (*112*). Both disciplines apply distinct methods that reflect these scale-related differences, and together they provide complementary views of plant functional diversity across scales. While field-based studies provide precise estimates of local functional diversity (α), remote sensing approaches are well suited to capturing broader-scale heterogeneity (β), especially when pixel size is comparable to the scale of plant communities.

Additionally, how functional diversity indices are calculated is often insufficiently reported in both disciplines, despite its importance in determining their ecological relevance. Field-based studies usually construct dissimilarity matrices from a small number of directly measured traits. In contrast, remote sensing studies often rely on high-dimensional spectral or modeled trait spaces, introducing additional preprocessing and dimensionality reduction steps. The construction of trait dissimilarity matrices –whether using Euclidean or Gower distances, or ordination techniques like PCA– can strongly influence the sensitivity and outcome of indices such as functional dispersion or Rao’s Q (*14*, *113*). These calculations usually require data preprocessing to standardize trait units and reduce trait redundancy (*14*, *22*). While these procedures are typically more direct in local, field-based studies, they can pose additional challenges in large-scale or remote sensing applications involving high-dimensional spectral data (particularly in approaches 2a and 2b). Without transparent reporting and harmonization of these computational steps, cross-study comparisons may reflect variation in analytical decisions as much as ecological differences (*51*). Nonetheless, recent advances have proposed methods to bridge the gaps between approaches. For instance, previous work introduced a generalizable normalization to make functional diversity metrics directly comparable independently of their dimensionality, which improved 1) the comparability of field-based and remotely-sensed functional diversity indices, and 2) the validation of plant functional diversity estimates from different sensors (*51*). This normalization can also be applied to compare functional diversity indices derived from field datasets that differ in the number or type of traits sampled. Together, these developments highlight that both disciplines are converging toward more interoperable and comparable methodological frameworks. The two main remote sensing approaches face distinct challenges that affect their capacity to provide ecologically significant estimates of functional diversity depending on how functional diversity is defined before remote sensing data is incorporated. Methods that integrate field-based traits and remote sensing enhance ecological inference. However, these methods must address spatial and temporal discrepancies between field and remote observations. Additionally, these methods must contend with trait variability driven by phenology, disturbances, or intraspecific phenotypic plasticity. Even traits that are strongly correlated at broad scales may respond differently to local environmental drivers, leading to scale-dependent shifts in correlations among traits (*114*). While this variability presents challenges in validating field-based and remote observations, it also presents an opportunity because remote sensing can capture these dynamics across larger spatial and temporal scales (*44*, *51*, *77*). However, remotely sensed data-only community-level methods, where functional diversity is inferred from spatial or spectral heterogeneity without defining trait spaces at the individual level, are limited by the difficulty of distinguishing vegetation-related variability from structural, background, and biochemical confounding effects in reflectance data, especially in heterogeneous landscapes. These limitations reflect the difficulty of translating pixel-level variability into meaningful ecological functional differences (*26*, *112*). In contrast, approaches that operate at the level of individual plants, where spectral variables can be attributed to individual plants, allow spectral information to estimate functional composition and diversity with similar techniques as used in field-based ecology [e.g., (*115*, *116*)]. While the quantitative relationship between spectral and functional diversity is still being refined, ongoing efforts are advancing the standardization and applicability of spectral diversity metrics for monitoring plant functional diversity (*45*, *112*). These methodological differences do not highlight shortcomings of any single approach, but rather, the need for careful validation to ensure conceptual and empirical alignment with field-based ecology.

In remote sensing, methodological differences in how abundance is incorporated into functional diversity indices primarily arise from how and at which ecological level diversity is defined before spectral information is incorporated. Based on this criterion, we identified two main remote sensing approaches: (1) integration with field-based data, where functional diversity is first quantified at the plot (or community) level using *in situ* trait measurements. Then, it is empirically linked to spectral or structural information using regression approaches [e.g., (*29*, *71*, *72*)]. Here, functional diversity is quantified independently of remote sensing, which is then used to predict (or extrapolate) these estimates across space; and (2) remotely sensed data-only methods. Two distinct scenarios arise within this category, depending on the size of pixels and plants. On one hand (approach 2a), when pixel resolution is coarser than individual plants, spectral variables represent a mixture of individuals and species within a pixel. Functional diversity is then inferred from spatial spectral heterogeneity or the diversity of traits estimated at such a scale, representing the CWM, and abundance is implicitly contained within the mixture of signals (*117*, *118*). In this case, although plant traits can be inferred from spectral data, they represent community-weighted trait means or mixed spectral signals within pixels rather than trait variation across individuals or species.

On the other hand (approach 2b), when pixel resolution or leaf spectra measurements [e.g., (*70*, *119*)], allows for the identification of individual plants or species, spectral variables can be attributed to them and assembled in a manner analogous to field-based ecology. First, traits or spectral features are assigned to individuals. Then, functional diversity indices are calculated by upscaling these per-plant estimates to the pixel level. This approach preserves the link between individual-level variation (i.e., intra- and interspecific trait variation) and community-level structure but requires high-resolution data and precise identification of individuals [e.g., (*120*)]. Together, these approaches demonstrate that the key distinction in remote sensing-based functional diversity is not whether plant traits are estimated, but rather how diversity is defined, and whether the spectral observations represent individual plants or aggregated species mixtures. This distinction further determines whether spectral observations represent individual plants or aggregated species mixtures, and how abundance is incorporated. When pixels integrate multiple individuals – or species – (approach 2a), abundance is implicitly embedded in the spectral signal, as reflectance contributions are weighted by the proportional cover or dominance of vegetation components within a pixel. In contrast, when individuals are resolved (approach 2b), abundance (e.g., biomass, cover, or frequency) can be integrated when calculating functional diversity indices, more closely mirroring field-based approaches. These differences influence how closely spectral variables approximate functional traits, and the degree to which remote sensing- and field-based estimates of functional diversity are comparable. When pixels are larger than plants, spectral mixtures dominate, increasing uncertainties due to window size and spatial resolution, and weakening trait-diversity relationships (*22*, *121*, *122*). Conversely, when individuals can be resolved, remotely-sensed spectral traits more closely approximate trait-based functional diversity [e.g., (*115*, *116*)]. Therefore, while pixel-based remote sensing is essential for large-scale, repeated monitoring, field-based methods provide the ecological precision needed to validate and establish trait estimates [(*38*, *41*); Fig. 4 and 5].

**Fig. 5.**
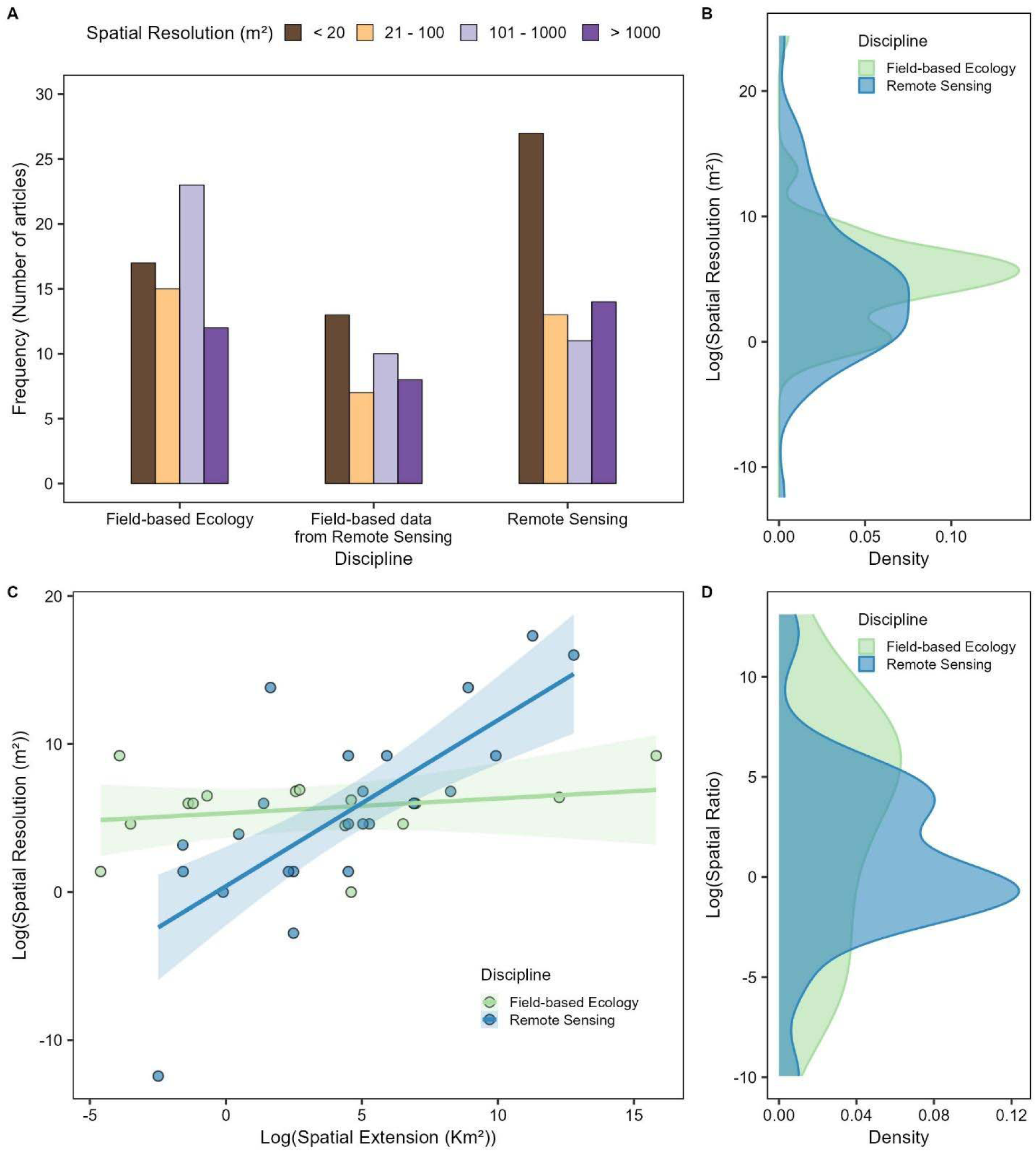
Spatial scales of functional diversity studies across disciplines. (**A**) Frequency of spatial resolutions evaluated in plant functional diversity research in field-based ecology, and remote sensing. Remote sensing is divided into field-based and exclusively remote sensed data. Field-based resolutions are plot areas, while in remote sensing resolutions are pixel areas. (**B**) Density of spatial resolution frequency (pixel size or plot size) by discipline. (**C**) Bi-variate relationship between spatial resolution and spatial extent for each discipline (**D**) Density of spatial ratio frequency by discipline. Spatial ratio is the ratio of spatial resolution to spatial extent and indicates the relative grain-to-extent relationship of each study. In (**B**), (**C**) and (**D**) we included only resolution and/or extent in remote sensing research.

### Spatial dimension and sensors

Our analysis reveals a systematic spatial decoupling between field-based ecology and remote sensing approaches to plant functional diversity (Fig. 5). Field-based studies focus on fine ecological scales, measuring traits at the individual- and/or organ-level and using plot sizes ranging from small (< 20 m^2^) to intermediate (101 - 1000 m^2^) sizes (Fig. 5A and B), which enables high-resolution characterization of trait variation within populations and communities. Although field-based studies can cover very large –but discontinuous– spatial extents (0.01 - 7,413,827 km^2^), remote sensing spans continuous spatial extents ranging from 0.01 to 358,000 km^2^. The resolution is defined by the size of the sensor pixels and ranges from individual plants in very high-resolution unmanned aerial vehicle (UAV) imagery to community or landscape-level representations in airborne and satellite platforms. Remote sensing exhibits a clear extent–resolution trade-off inherent to sensor design, but field-based studies do not show the same trade-off (Fig. 5C and D). Increasing the spatial extent of field surveys often comes at the cost of reduced replication or spatial representativeness, which can lower the precision of functional diversity estimates at broader scales.

Previous work across both field-based ecology and remote sensing has shown that ecological patterns are driven by processes operating at multiple spatial and temporal scales, and there is no ideal scale for capturing them (*24*). For example, a grassland study demonstrated that spectral diversity is highly sensitive to the resolution of observation, with distinct patterns emerging across five spatial scales (*70*). Additionally, simulation- and aggregation-based studies have shown that field data aggregated to coarser pixel sizes can help recover trait-spectrum linkages when pixels integrate multiple species (*21*, *22*). Similarly, biodiversity-ecosystem functioning relationships have been shown to be nonlinear across spatial scales, with increasing community aggregation and complexity at broader extents (*123*). We found that as remote sensing increases the spatial extent of the study area the spatial resolution decreases (Fig. 5C and D), and reflecting an inherent trade-off in satellite mission design that can be partially overcome by combining data of multiple images [e.g., (*32*)]. In contrast, field-based studies cover very large areas, but they use spatially discrete sampling to represent large spatial extents rather than spatially continuous sampling. Still, the use of coarse spatial resolutions could limit the capacity to capture fine-scale ecological variation or to effectively scale up patterns to larger extents, while sparse field sampling may miss spatial heterogeneity occurring between sampling units. To better understand how ecological processes shape functional diversity patterns, we need to explicitly integrate the hierarchy of spatial scales because ecological processes operate in a scale dependent manner (*24*, *123*). Emerging frameworks such as spectral biology (*124*), offer a promising way to connect spectral data with plant functional diversity patterns across spatial scales or temporal scales.

Scale-related trade-offs underscore the essential role of field-based trait data in remote sensing applications, as remote sensing studies rely on field-based information to calibrate and/or validate trait estimates (Fig. 6A). Remote sensing alone cannot capture all functional traits, particularly those lacking reliable optical signatures [e.g., root traits, seed mass, and floral traits; (*24*)]. Although very high-resolution UAV and commercial satellite imagery can distinguish individual plants in certain ecosystems, field measurements continue to provide the fine ecological resolution and direct trait information that remote sensing cannot yet capture consistently. These field data enable calibration, validation, or trait retrieval. Indeed, 80% of remote sensing studies incorporated field-based information from literature, databases, and/or *in situ* measurements (Fig. 6A). This integration is crucial to ensure that remotely-sensed functional diversity reflects ecological reality by linking fine-scale processes with broader-scale patterns.

**Fig. 6.**
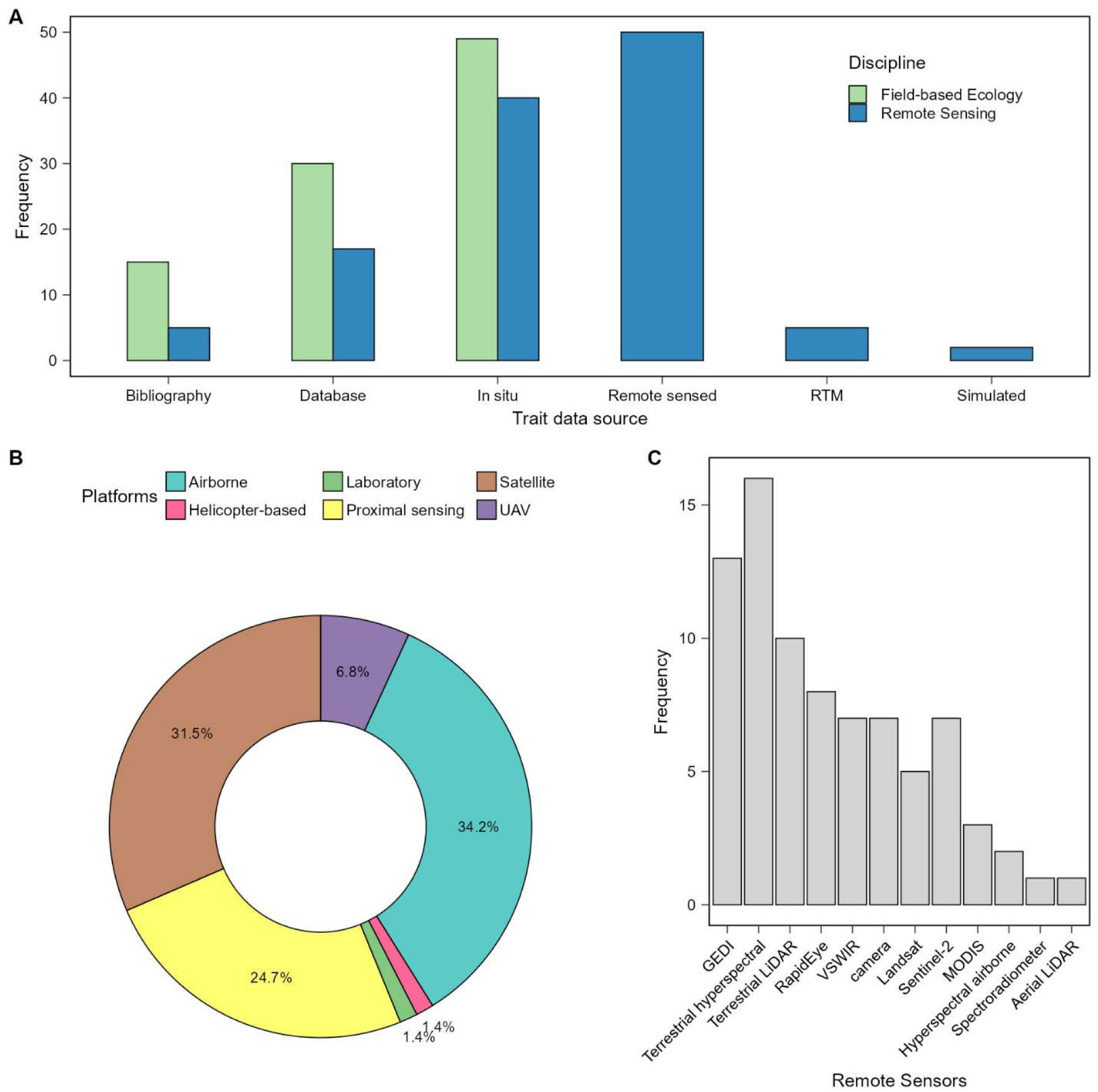
Sources and platforms of plant trait data. (**A**) Frequency of trait data sources used in field-based ecology and remote sensing research. (**B**) Frequency of remote sensing platforms used to retrieve plant traits. (**C**) Frequency of remote sensors employed in the reviewed studies. UAV: Unmanned aerial vehicle.

The use of satellite, airborne, and proximal sensing platforms among sensors is now relatively balanced for evaluating plant functional diversity patterns (Fig. 6B and C), reflecting the growing availability and accessibility of diverse remote sensing platforms and sensors. Satellite-based platforms remain widely used due to their global coverage, regular revisit intervals, and increasing data availability. Their role is expected to grow as new hyperspectral missions become operational (e.g., EnMAP, PRISMA, CHIME, and/or HISUI). However, their moderate spectral and spatial resolutions limit their utility for detection of biochemical and structural traits (*24*, *123*). Hyperspectral sensors, especially those on airborne or UAV platforms, offer higher spectral fidelity that can resolve fine-scale trait variation, albeit at the cost of spatial extent (*46*, *120*). Overcoming the limitations of individual platforms or resolutions requires explicit multiscale frameworks (*109*) that can link local trait variation to broader functional diversity gradients across landscapes. Therefore, integrating field-based and remote sensing data within multiscale frameworks is particularly promising for capturing both local trait expression and broad ecological gradients, underscoring that these platforms are most powerful when used together and not separately.

### Future directions and outstanding questions

Despite recent progress, our review reveals important conceptual and methodological gaps that still limit a fully integrated cross-scale understanding of plant functional diversity. These gaps do not reflect the shortcomings of either discipline. Instead, they arise from the uneven distribution of research efforts across ecosystems and data types. We found that an urgent priority is to expand trait-based research into biomes that are largely absent from the literature [e.g., deserts, alpine systems, palustrine wetlands, and other land-water interface biomes; (*69*, *125*)]. These ecosystems are often difficult to access and less suitable for many satellite and airborne sensor systems [e.g., (*60*, *62*)], yet they are critical for understanding how biodiversity responds to global change. This raises the question of how we can systematically extend the monitoring of functional diversity monitoring into ecosystems with low vegetation cover or high environmental heterogeneity, using either field-based or remote sensing approaches, or both. Addressing this issue will require targeted sampling campaigns, novel sensor configurations, and adaptive analytical frameworks that can accommodate sparse or patchy vegetation signals (*126–128*).

It is equally urgent to harmonize the selection, measurement, and definition of traits across disciplines. We found that field-based ecology tends to measure structural or reproductive traits, whereas remote sensing focuses on biochemical and spectral traits, limiting interoperability and undermining cross-scale analysis (*103*, *129*). Advancing integration requires identifying a core set of traits that can be consistently measured and meaningfully compared across both disciplines, and at what spatial resolutions? As highlighted in spatial scaling theory (*24*), there is no universally optimal resolution. Thus, the ecological meaning of a resolution depends on plant size, density, and spatial configuration rather than on its absolute value. Recent modelling and simulation-based work is beginning to formalize operational resolution limits beyond which remotely sensed estimates of functional diversity lose ecological interpretability (*130*). More broadly, spatial resolution has been shown to critically influence the interpretation of functional diversity and community assembly processes (*110*, *131*), reinforcing the need to explicitly account for the relationship between pixel size, plant size, and community structure when designing biodiversity monitoring frameworks. Progress towards this will depend on developing community-driven standards for trait definitions and metadata, as well as trait weighting, dimensionality reduction, and the incorporation of intraspecific variation (*15*, *108*, *132*).

Methodological integration demands the systematic incorporation of multiscale, spatially explicit frameworks (*21*, *50*) that can link fine-scale ecological processes with broader landscape and regional patterns (*56*, *123*, *133*). Recent advances demonstrate that regional-to-continental modelling of canopy traits is feasible when multiscale data are combined with appropriate analytical frameworks (*134*). Unresolved issue concerns which monitoring designs best capture the interplay between local trait variation and regional biodiversity patterns. Bridging the scale gap requires multi-tiered approaches combining field plots, airborne campaigns, satellite platforms, and coordinated sampling designs that ensure that trait measurements, spectral data, and structural information are collected at compatible spatial and temporal scales (*31*, *135*). This reconciles spatially discrete field sampling across large areas and spatially continuous remote sensing. Rather than emphasizing disciplinary limitations, this integrative perspective highlights that each discipline samples different portions of the ecological scale continuum. Field-based ecology provides fine-resolution, mechanistic trait information, whereas remote sensing captures broad spatial and temporal patterns that can not be achieved through ground-based sampling alone. Emerging very high-resolution UAV imagery increasingly bridges this gap by matching individual-level ecological scales. Thus, advancing plant functional diversity research will depend not on privileging one discipline over the other, but on strategically integrating both within coordinated multiscale frameworks (*111*, *136*).

Finally, the robustness of using remote sensing to predict and evaluate functional diversity rests on empirically validating spectral proxies and derived metrics with field measurements (*29*). While such validation efforts exist, they are uneven across biomes and geographic regions, particularly in ecosystems where spectral signals are sparse, noisy, or strongly confounded by background effects (*137*, *138*). Notably, large coordinated initiatives such as National Ecological Observatory Network (NEON) demonstrate that systematic, plot-level validation across multiple sites is feasible when field and airborne observations are jointly designed (*100*). This underscores the extent to which spectral proxies may be generalized across ecosystems, and conditions under which site- or biome-specific calibration remain necessary. Strengthening these empirical links will enhance the predictive power of trait-based approaches and facilitate their integration into biodiversity conservation and monitoring programs (*139*, *140*). Ultimately, fostering deeper conceptual and methodological convergence between field-based and remote sensing will help close remaining methodological gaps and address the overarching question.

## Materials and Methods

### Data collection

We developed this review following standard roadmaps for literature reviews (*141*), and we modified the Reporting standards for systematic evidence syntheses (ROSES) Flow Diagram for Systematic Maps and Systematic Review (Fig. S1) (*142*). We explored published literature on plant functional diversity from 1976 to the end of 2024 by searching for relevant documents in the Web of Science (WoS) and the Scopus databases for relevant documents. We conducted the search in April 2025 by designing a search query with keywords to identify literature related to plant functional diversity (i.e., *trait composition*, *functional composition*, *functional richness*, *trait diversity*, and *functional diversity*). These terms represent widely used descriptors of functional diversity in plant ecology, as established in foundational literature (*10*, *39*). First, we added exclusion criteria related to ecosystems and specific subject areas to constrain the query to terrestrial, natural applications (Supplementary materials). We harmonized subject areas as they differ between databases (e.g., Agricultural and Biological Sciences (“AGRI”) from Scopus and Agriculture Multidisciplinary (or Agronomy) from WoS). Then, we divided the literature into field-based ecology and remote sensing (see subsection *Data analysis*). We initially obtained 10,955 records from WoS and 7,594 records from Scopus (Fig. S1). We then eliminated duplicate documents based on their Digital Object Identifier (DOI) or title. Finally, we filtered by document type, keeping articles, conference papers and data papers and excluding editorial papers, corrections, meeting abstracts, short surveys, notes and reviews. We retained a total of 11,243 original research articles (Fig. S1). We restricted the search to English-language literature to ensure terminological consistency and reproducibility across databases.

### Data analysis

We identified the main topics and trends of the selected literature using a combination of bibliometric and scientometric analyses. We divided the selected literature to compare plant trait diversity research based on field ecology and remote sensing (Fig. S1) using methodological terms associated with each in the title, abstract, and keywords (author’s keywords and keywords associated with the database) (see Supplemental material for the list of terms). We included studies measuring in natural, semi-natural, and experimental fields as field-based ecology.

### Bibliometric analysis

We performed a comprehensive bibliometric analysis of the selected literature using the R package “bibliometrix” (*143*). We estimated: 1) productivity by discipline, including timespan, amount of different sources and documents, annual growth rate, document mean age, mean citations per document, mean citations per document per year, and number of references; and 2) proportion of keywords per year by discipline, including the number of published articles in which a keyword was used over the total number of documents in the database of the respective discipline. We used authors’ keywords and keywords provided by WoS and SCOPUS. We carried out both analyses with the *biblioAnalysis* function.

We built a keyword co-occurrence network for each discipline with the *biblioNetwork* function, connecting each pair of the 30 most common keywords and estimated the proximity between keywords (modularity) and the frequency of each keyword. We used the indicator *K* –the ratio between the average number of unique keywords and the mean keyword frequency of each discipline (*144*)– to assess the scientific maturity of each discipline. The indicator assumes that as a discipline matures conceptual consensus is reached, which reduces the relative frequency of unique keywords (analogous to concepts). We estimated this indicator for each year to capture temporal trends for each discipline, as the ratio between keyword count and keyword frequency. *K* values between 1 and 0.75 indicate a pre-revolution or revolution stage of a discipline (immature), between 0.75 and 0.5 is a forming or pre-normal science stage, between 0.5 and 0.25 a pre-normal or normal science stage, and between 0.25 and 0 is a post-normal science or next pre-revolution stage (mature).

### Structural Topic Model

We fitted a structural topic model (STM) with the “quanteda” R package (*145*) to identify and characterize the underlying topics of the selected publications based on their title, keywords, and abstracts. This model allowed us to explore key topics in the scientific literature on plant functional diversity, their temporal patterns, and differences in their relative importance between disciplines (field-based ecology and remote sensing, as classified in subsection *Data analysis*). First, we identified the optimal number of topics in which the articles were grouped with the *stm* function from the “stm” R package. We tested several models with different numbers of topics between 2 and 10, as the analysed texts are not extensive, and the query is not too specific (*146*). We selected the model with seven topics, as it best represented the tradeoff between semantic coherence and exclusivity. This choice was supported by model diagnostics (held-out likelihood, lower bound, residuals, and the coherence-exclusivity balance), which indicated that seven topics provided the most stable and interpretable solution (Fig. S2). We reviewed and ranked the probability of each topic that an article belongs to (gamma, γ). Then, we assigned each article to a topic based on the highest gamma and calculated the annual proportion of articles per topic. We identified the topics based on the complete plant functional diversity articles database. In addition, we tested the effects of discipline on the relative importance of the identified topics with the *estimateEffect* function from the “stm” R package.

### Inclusion criteria for the systematic review

Due to the large amount of articles obtained in our literature search, we designed criteria to select the most impactful articles in the scientific literature on plant functional diversity (*141*). To evaluate temporal trends, we divided the literature into four eight-year cohorts (2000–2024) and selected the most-cited articles within each cohort. Because the volume of publication differed substantially between disciplines, we applied different thresholds: the 25% most-cited papers in remote sensing (n = 142) and the 2.5% most-cited papers in field-based ecology (n = 268; Fig. S1). These unequal thresholds reflect the strong disparity in the number of documents between the disciplines, and were chosen to obtain comparable sample sizes rather than equivalent percentiles. We reviewed the abstracts of all selected articles and retained 42.25% (n = 60) of the remote sensing literature and 35.45% (n = 95) of the field-based ecology literature (Fig. S1). We further excluded articles that were unrelated to vegetation or that studied diversity but not functional diversity (e.g., only taxonomic and/or phylogenetic diversity). We also excluded articles that studied vegetation but measured functional traits in a taxonomic group other than plants. We further excluded articles on new conceptual frameworks and reviews because they did not directly measure plant functional diversity, but we included those that performed meta-analyses in which functional diversity was analysed. Finally, we included a total of 56 remote sensing and 74 field-based ecology articles in the systematic review (Fig. S1). From each article, we extracted the following information from the selected articles: spatial grain and extension, ecological aspects (e.g., biomes, traits, and species’ abundances), methodological approaches (e.g., how is trait information collected, and remote sensing platforms and sensors), and statistical aspects (e.g., functional diversity indices, and whether they were weighted by abundance; further details are provided in Table S1). We standardized the names of plant traits and grouped them based on the AusTraits Plant Dictionary (APD) (*147*). Also, we assigned biomes under study according to function-based classification of ecosystems (*148*). We performed all analyses in R v. 4.6.0 (*149*).

## Acknowledgments

We thank Anna K. Schweiger for helpful comments, constructive suggestions, and valuable perspective.

## Funding

Data Observatory Foundation PhD fellowship (JMC-P).

Agencia Nacional de Investigación y Desarrollo (ANID), Fondecyt N° 11241088 (JL).

“Integrated Observing Systems and Simulation Experiments to Analyze Biodiversity-Ecosystem Function Relationships in Savanna Ecosystems” PID2023-151046NB-I00 funded by MCIU/ AEI / 10.13039/501100011033 / FEDER, UE) (JP-L).

## Author contributions

Conceptualization: JMC-P, DC, JL

Methodology: JMC-P, DC, JL

Formal analysis: JMC-P

Visualization: JMC-P

Data Curation: JMC-P

Supervision: DC, JL

Writing—original draft: JMC-P, DC, JL

Writing—review & editing: JP-L, DC, JL

## Data and materials availability

All relevant results are present in the main text and the Supplementary Materials. The code and data used in this study are available at the following Zenodo repository: https://doi.org/10.5281/zenodo.18684166.

