## Supplementary materials for "Bridging the gaps between field-based ecology and remote sensing to estimate plant functional diversity: a systematic review"

José M. Cerda-Paredes *et al.*

**This PDF file includes:**

Supplementary Text

Figs. S1 to S7

Tables S1 to S3

References (*142*, *147*, *148*)

**Supplementary Text**

Queries

WoS

("trait composition" OR "functional composition*" OR "functional richness*" OR "trait diversit*" OR "functional diversit*") AND ("plant" OR "tundra" OR "taiga" OR "desert" OR "savanna" OR "forest" OR "grass*" OR "wetland*" OR "mangrove*" OR "shrubland*" OR "scrubland*" OR "tree" OR "shrub" OR "herb*") (All Fields) and Ecology or Plant Sciences or Environmental Sciences or Biodiversity Conservation or Forestry or Soil Science or Multidisciplinary Sciences or Microbiology or Evolutionary Biology or Agronomy or Marine Freshwater Biology or Biochemistry Molecular Biology or Geography Physical or Biology or Biotechnology Applied Microbiology or Agriculture Multidisciplinary or Genetics Heredity or Geosciences Multidisciplinary or Zoology or Environmental Studies or Cell Biology or Water Resources or Engineering Environmental or Mycology or Urban Studies or Green Sustainable Science Technology or Imaging Science Photographic Technology or Chemistry Multidisciplinary or Horticulture or Meteorology Atmospheric Sciences or Food Science Technology or Biochemical Research Methods or Biophysics or Geography or Regional Urban Planning or Mathematical Computational Biology or Energy Fuels or Agricultural Engineering or Physiology or Virology or Anatomy Morphology or Computer Science Interdisciplinary Applications or Developmental Biology or Statistics Probability or Computer Science Information Systems or Computer Science Theory Methods or Mathematics Applied or Paleontology or Remote Sensing or Geochemistry Geophysics or Architecture or Archaeology or Computer Science Artificial Intelligence or Development Studies or Spectroscopy or Oceanography or Limnology or Entomology (Web of Science Categories)

Scopus

TITLE-ABS-KEY ( ( "trait composition" OR "functional composition*" OR "functional richness*" OR "trait diversit*" OR "functional diversit*" ) AND ( "plant" OR "tundra" OR "taiga" OR "desert" OR "savanna" OR "forest" OR "grass*" OR "wetland*" OR "mangrove*" OR "shrubland*" OR "scrubland*" OR "tree" OR "shrub" OR "herb*" ) ) AND ( INCLUDE ( SUBJAREA , "AGRI" ) OR INCLUDE ( SUBJAREA , "COMP" ) OR INCLUDE ( SUBJAREA , "EART" ) OR INCLUDE ( SUBJAREA , "ENGI" ) OR INCLUDE ( SUBJAREA , "ENVI" ) OR INCLUDE ( SUBJAREA , "IMMU" ) OR INCLUDE ( SUBJAREA , "MATH" ) OR INCLUDE ( SUBJAREA , "MULT" ) )

Discipline filter

(remot*, sensin*, teledetec*, Sentinel*, Landsat*, LiDAR*, satellit*, MODIS*, spectroscop*, aerial*, dron*, hyperspect*, UAV, UAS o RPAS)

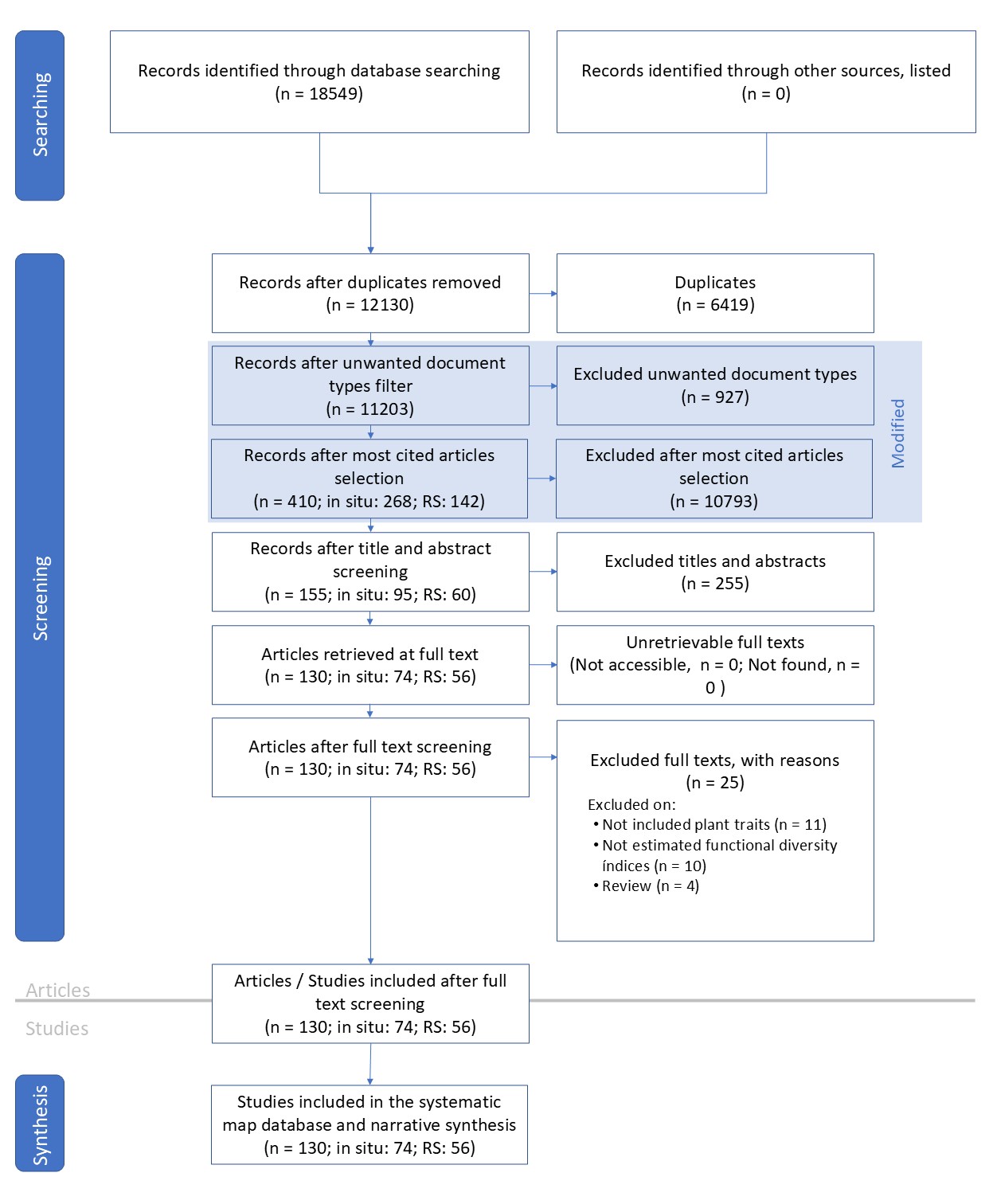

**Fig. S1.**

Explanation diagram of the roadmap of the review. Combinated ROSES flow diagrams for systematic maps and review (*142*).

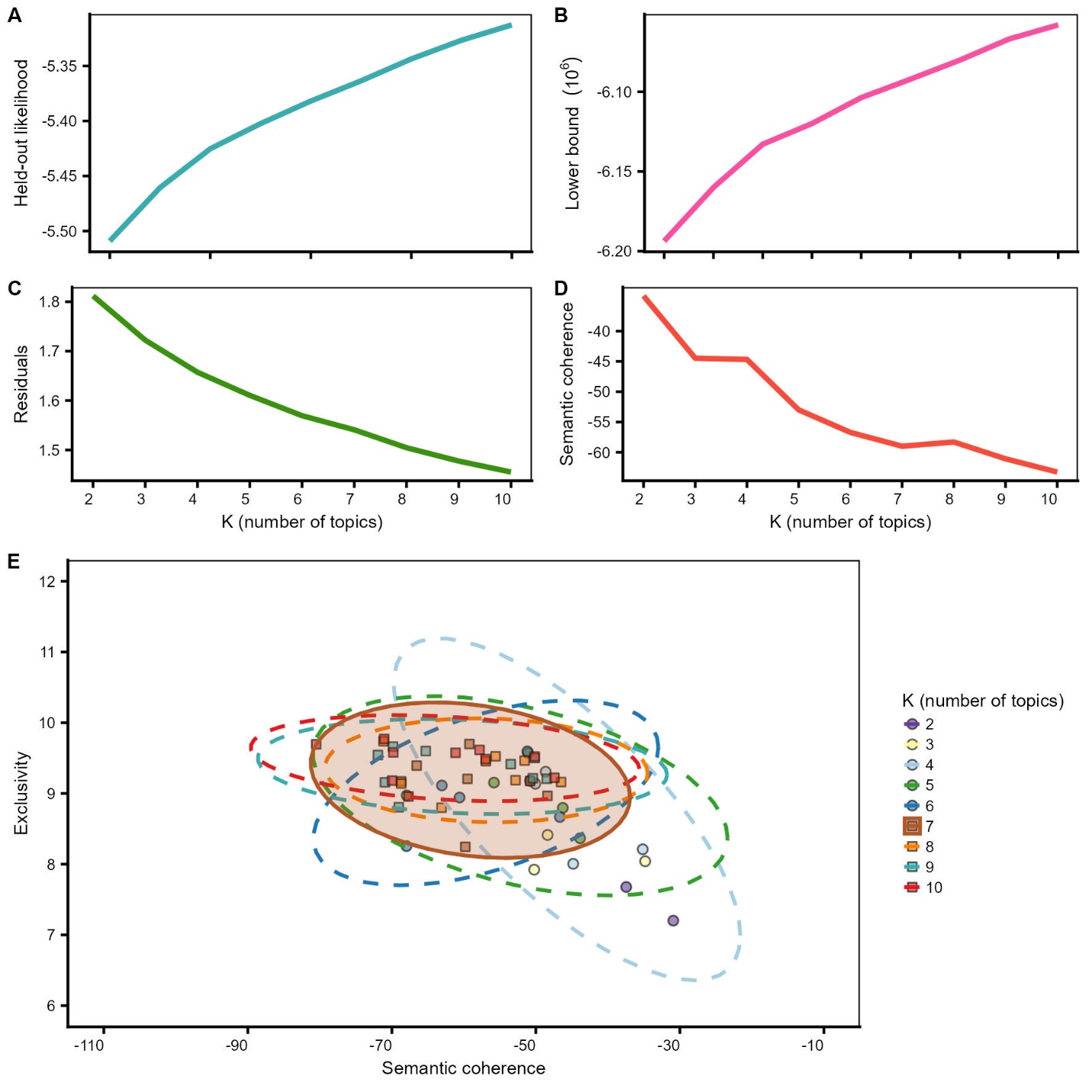

**Fig. S2.**

Model selection diagnostics for the structural topic model (STM) across different numbers of topics (K = 2–10). **a)** held-out likelihood. **b)** lower bound of the log-likelihood. **c)** residuals. **d)** semantic coherence. **e)** Trade-off between semantic coherence and topic exclusivity for each K value, with ellipses representing the dispersion of topics at a given K value. The shaded ellipse is K = 7, the selected model.

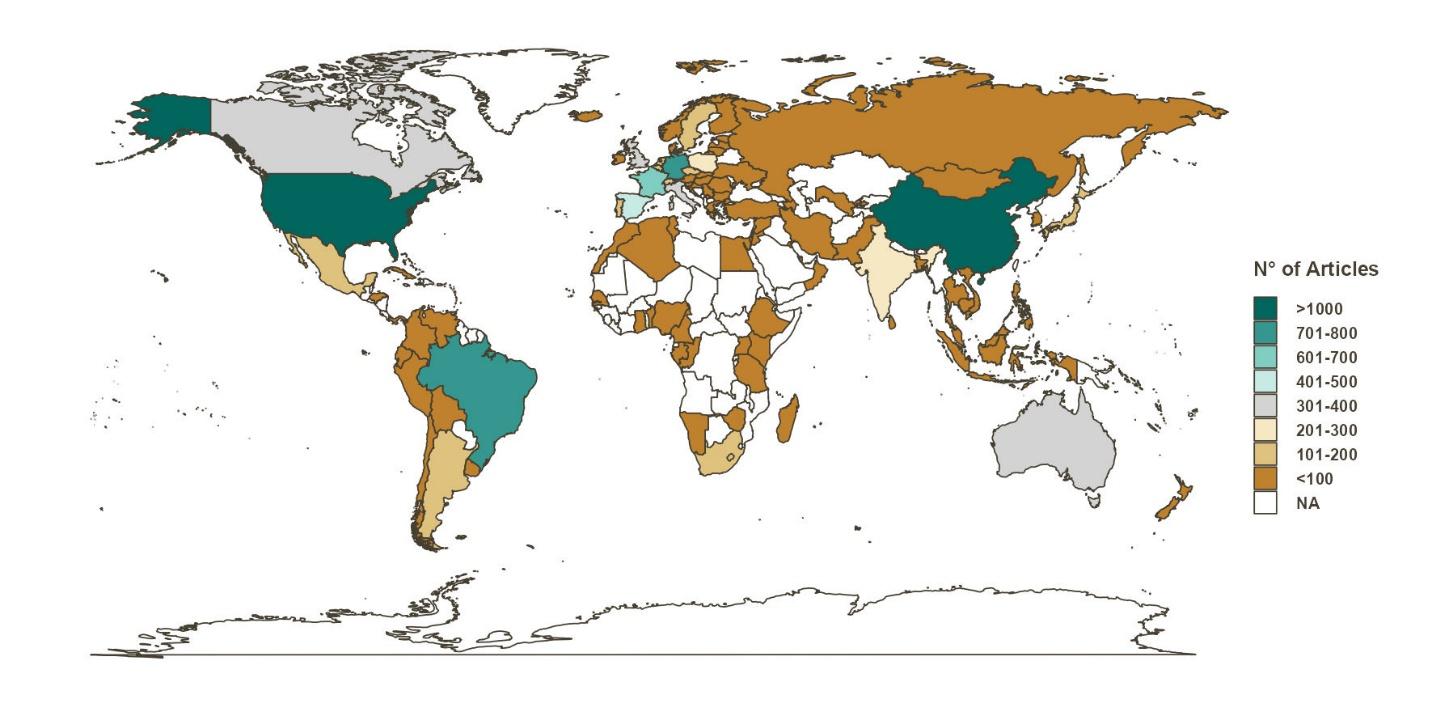

**Fig. S3.**

Number of articles published by countries on plant functional diversity. The countries depict the corresponding author’s affiliation.

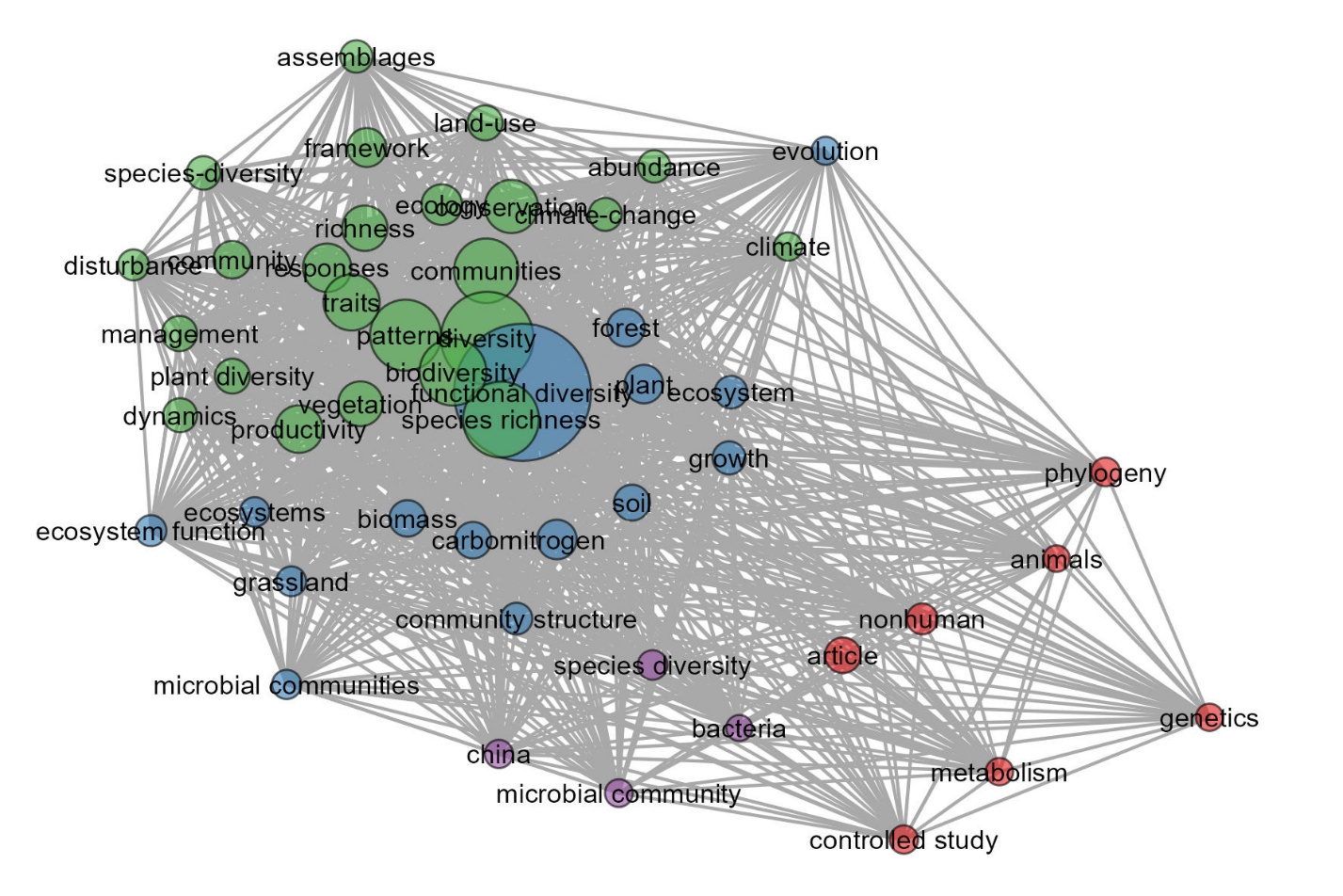

**Fig. S4.**

Network keyword co-occurrence in field-based ecology of the top 50 most used keywords. The size of the nodes represents the frequency of the keyword in the network, and the distance between them indicates the co-occurrence frequency between the keywords. The width of the links represents the strength of the connection (i.e., the co-occurrence). The colors of the nodes represent their clusters.
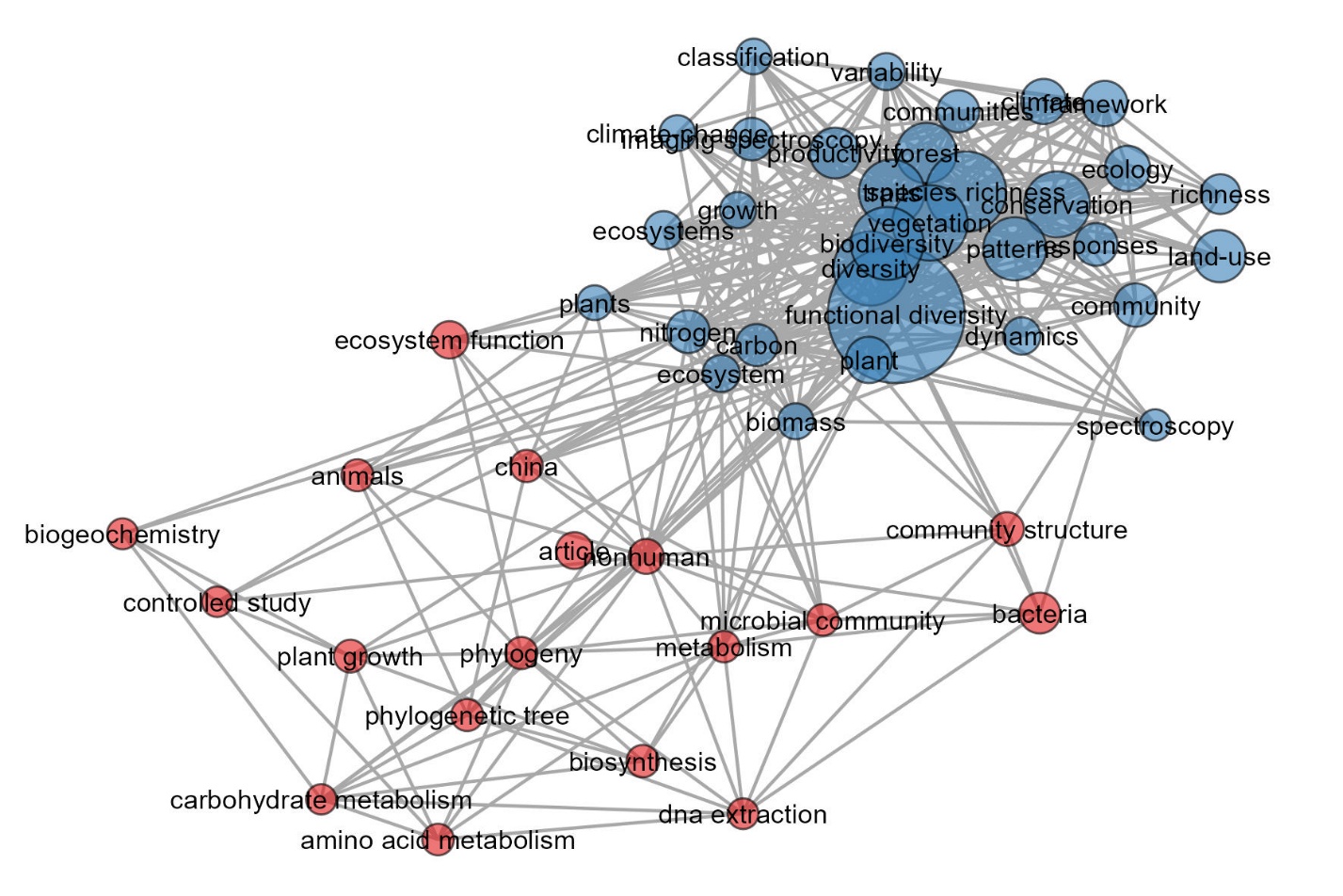

**Fig. S5.**

Network keyword co-occurrence in remote sensing of the top 50 most used keywords. The size of the nodes represents the frequency of the keyword in the network, and the distance between them indicates the co-occurrence frequency between the keywords. The width of the links represents the strength of the connection (i.e., the co-occurrence). The colors of the nodes represent their clusters.

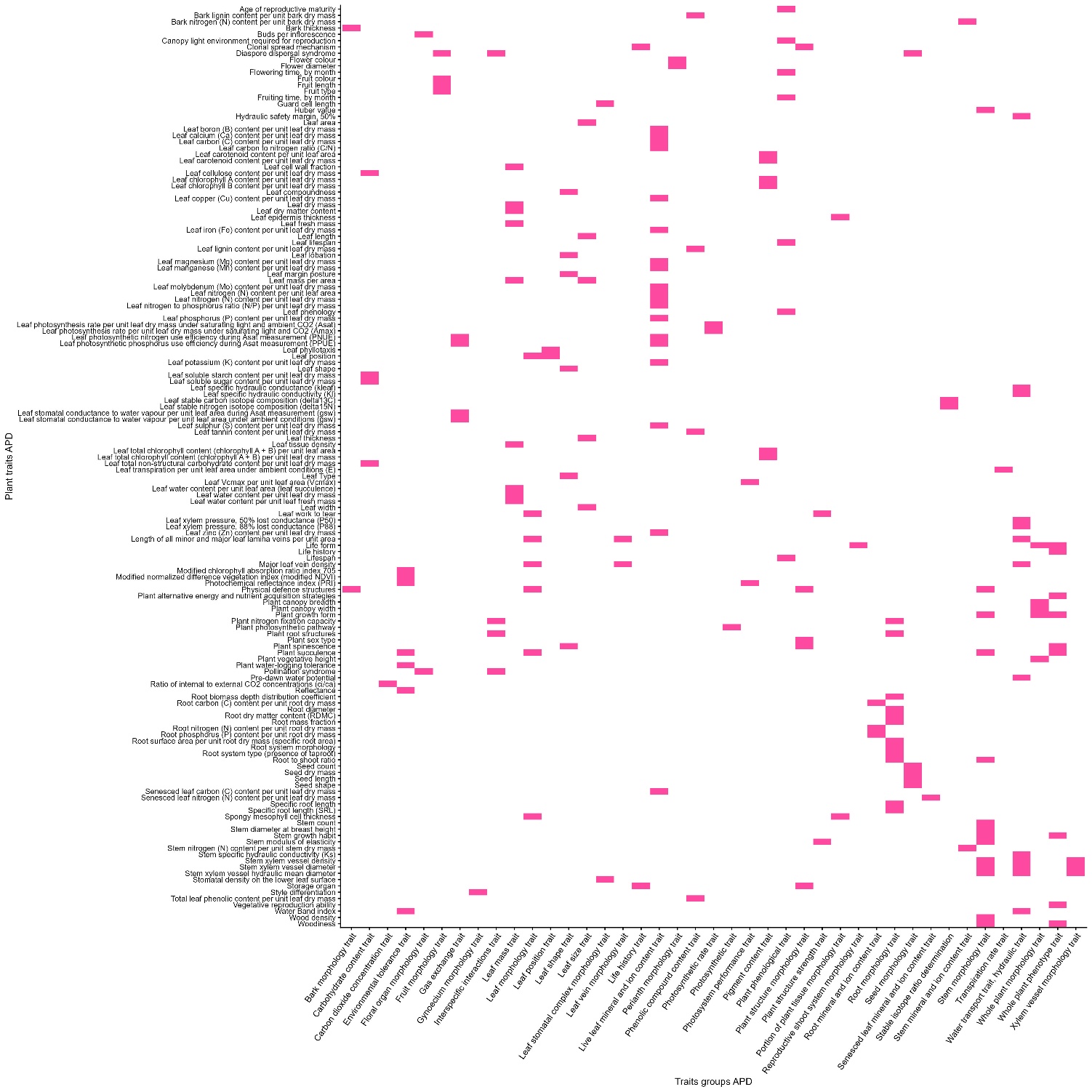

**Fig. S6.**

Representation of each plant trait registered to groups of traits based on AusTraits Plant Dictionary (APD) (*147*).

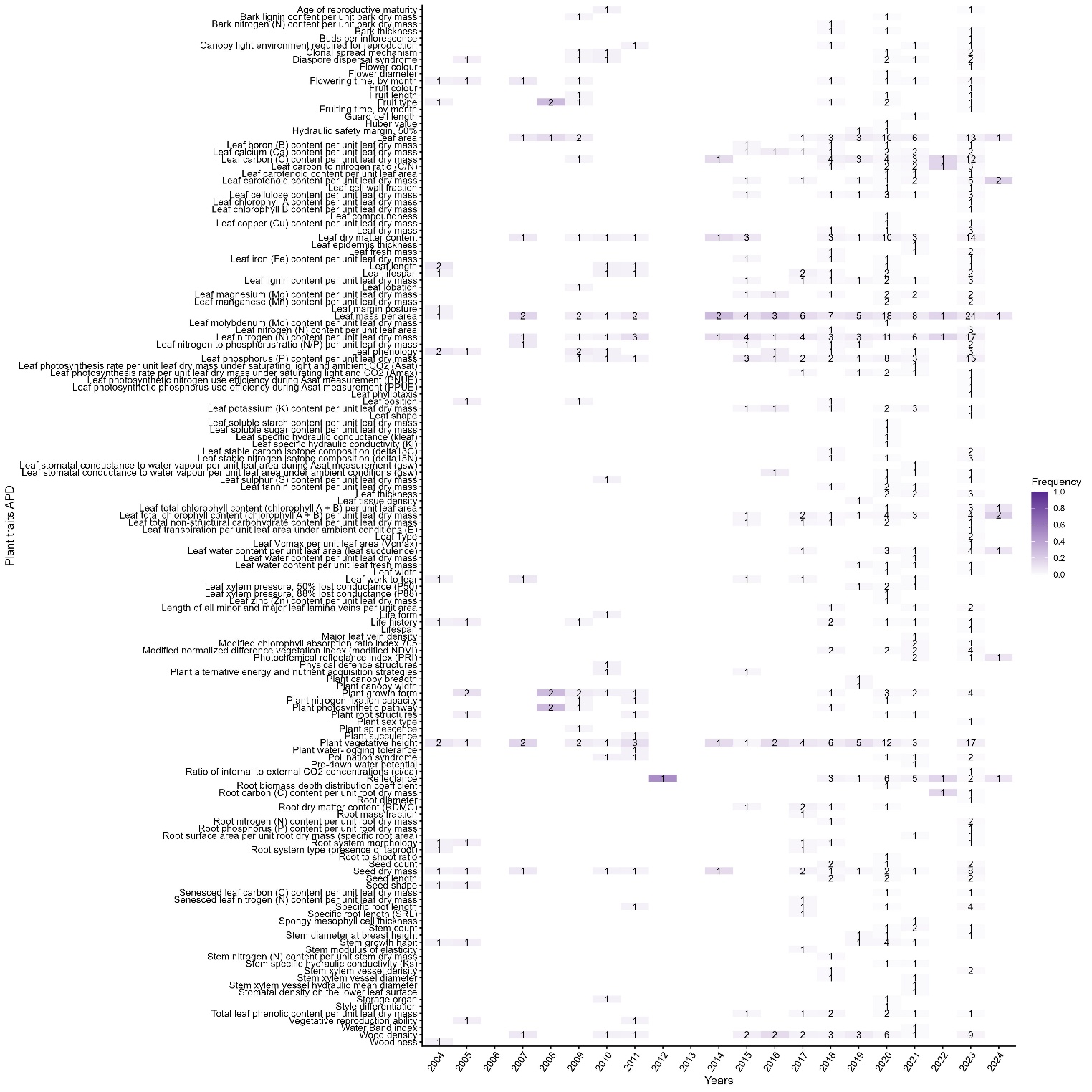

**Fig. S7.**

Heatmap of plant traits identified in each biome per discipline. Cell color represents the proportion of each trait per biome, and cell numbers represent the number of articles.

**Table S1.**

Systematic review parameters.

| **parameter** | **details** |
| --- | --- |
| spatial_extension | spatial extension (km^2^) |
| spatial_resolution | spatial resolution; pixel size for remote sensors or minimum sampling unit size in field work (m^2^) |
| biome | biome under study (*148*) |
| platform | platform where the sensor is located |
| sensors | sensor's name |
| trait_def | traits involved in the study (*147*) |
| group_def | groups to which the traits included in the study belong (*147*) |
| diversity_index | metrics applied to traits |
| abundance | Is the species abundance considered in the study? (BOOLEAN, yes = 1, no = 0, not specified = empty) |
| abundance_type | how traits are weighted if they are |
| sampling_methodology | if "abundance" is TRUE, how did they sample? |
| trait_data_origin | origin of the data, whether from field work, literature, databases, remotely sensed |

**Table S2.**

Summary production per discipline

| **Description** | **Field-based Ecology** | **Remote Sensing** |
| --- | --- | --- |
| Timespan | 1976–2024 | 2002–2024 |
| Sources (Journals, Books, etc) | 1196 | 240 |
| Documents | 10659 | 544 |
| Annual Growth Rate % | 15.89 | 22.18 |
| Document Average Age | 6.99 | 5.79 |
| Average citations per doc | 27.32 | 20.49 |
| Average citations per year per doc | 2.544 | 2.32 |
| References | 347223 | 31344 |

**Table S3.**

Estimated effect of disciplines on STM topics.

| Topic | Coefficient | Estimate | SE | t | p |
| --- | --- | --- | --- | --- | --- |
| 1 | (Intercept) | 0.102 | 0.002 | 48.987 | 0.000 |
|  | discipline | -0.010 | 0.009 | -1.046 | 0.296 |
| 2 | (Intercept) | 0.222 | 0.002 | 89.326 | 0.000 |
|  | discipline | -0.004 | 0.011 | -0.345 | 0.730 |
| 3 | (Intercept) | 0.136 | 0.002 | 60.130 | 0.000 |
|  | discipline | -0.049 | 0.010 | -5.063 | **0.000** |
| 4 | (Intercept) | 0.180 | 0.002 | 85.480 | 0.000 |
|  | discipline | -0.009 | 0.009 | -1.010 | 0.313 |
| 5 | (Intercept) | 0.144 | 0.002 | 86.850 | 0.000 |
|  | discipline | 0.101 | 0.009 | 11.280 | **0.000** |
| 6 | (Intercept) | 0.121 | 0.002 | 71.757 | 0.000 |
|  | discipline | -0.008 | 0.008 | -1.052 | 0.293 |
| 7 | (Intercept) | 0.095 | 0.001 | 68.809 | 0.000 |
|  | discipline | -0.020 | 0.006 | -3.076 | **0.002** |
